# Antibody co-administration robustly improves proton therapy with radiosensitizing nanoparticles: a mathematical modeling study

**DOI:** 10.64898/2026.08.30.748121

**Authors:** Maxim Kuznetsov, Andrey Kolobov

**Affiliations:** Mathematical Oncology Laboratory (MOLAB), Institute of Applied Mathematics in Science and Engineering (IMACI), University of Castilla-La Mancha, Avda. Camilo José Cela 3, Ciudad Real, 13071, Spain; Division of Theoretical Physics, P.N. Lebedev Physical Institute of the Russian Academy of Sciences, 53 Leninskiy Prospekt, Moscow, 119991, Russia

**Keywords:** binding-site barrier, enhanced permeability and retention, receptor competition, intratumoral drug delivery, virtual cohort

## Abstract

Radiosensitizing nanoparticles represent a promising approach for enhancing the efficacy of proton radiotherapy; however, their performance is constrained by restricted penetration into tumor tissue, resulting in preferential perivascular accumulation. Here, we develop a spatially distributed mathematical model of a growing tumor undergoing proton therapy with intravenously administered radiosensitizing nanoparticles to investigate treatment optimization strategies. Using physiologically plausible parameter ranges informed by our own experimental measurements and published data, we demonstrate that co-administration of targeted nanoparticles with antibodies binding to the same tumor receptors can overcome transport-induced localization and promote a more uniform intratumoral redistribution of nanoparticles before irradiation. Population-level simulations across heterogeneous parameter sets suggest that moderate antibody doses consistently prolong tumor regrowth time, whereas higher antibody doses produce a pronounced and robust increase in tumor cure probability under a single high-dose irradiation regimen representative of preclinical settings.

A key conceptual result of our analysis is the asymmetric risk associated with antibody co-administration. In contrast to antibody–drug conjugates, for which excessive dosing of unconjugated antibodies may severely compromise therapeutic efficacy, co-administration of antibodies with nanoparticle-based radiosensitizers constitutes a “safe-by-design” strategy with respect to tumor cell kill in the modeled single high-dose irradiation setting: although excessive antibody doses may yield suboptimal outcomes, they cannot reduce tumor cell kill below that achieved with targeted nanoparticles administered without antibodies. These findings identify antibody-mediated spatial redistribution of radiosensitizing nanoparticles as a favorable strategy that is expected to provide robust therapeutic benefit despite substantial variability in tumor characteristics.

## 1. Introduction

In recent years, there has been a growing interest in the use of non-radioactive elements in proton therapy, both for improving tumor tissue imaging [1] and for enhancing the efficacy of radiation treatment through local radiosensitization of tumor cells [2–4]. Radiosensitizing systems are typically based on nanoparticles containing boron-11 or high-atomic-number elements, stabilized by biocompatible polymers and optionally functionalized with tumor-specific ligands [5–8]. Different chemical elements exhibit distinct mechanisms of radiobiological enhancement; for example, proton irradiation of gold leads to the emission of secondary and Auger electrons and induces the formation of hydroxyl radicals in the surrounding aqueous medium [9].

The effectiveness of radiosensitizing nanoparticles in proton irradiation has been confirmed by numerous experimental studies. In *in vitro* models, an increase in the frequency of DNA strand breaks [10], disruption of proliferation, and induction of apoptosis in tumor cells [11, 12] have been demonstrated, consistently leading to a substantial reduction in cell survival [13]. For example, in the experiments by Cunningham et al. [10], the use of gold nanoparticles resulted in an increase in ovarian cancer cell killing by ≈27% at an irradiation dose of 2 Gy and by ≈44% at a dose of 6 Gy. In the work of our experimental group, gold nanoparticles at concentrations of ≥25 µg/mL enabled complete suppression of the clonogenic activity of Ehrlich carcinoma cells at an irradiation dose of 4 Gy [14].

The results of *in vivo* experiments using mouse models demonstrate the ability of radiosensitizing nanoparticles to provide enhanced tumor growth suppression compared to proton irradiation alone. However, the corresponding improvement in therapeutic efficacy is generally less pronounced than that observed in *in vitro* experiments [14–16]. One of the key barriers limiting the effectiveness of nanoparticles in solid tumors is the restricted delivery of the agent into tumor tissue through the capillary network. This fundamental challenge is well known for tumor-specific antibodies with a hydrodynamic radius of 5–6 nm, for which typically only a small fraction of the intravenously administered dose reaches the tumor [17]. The problem becomes even more severe for nanoparticles with sizes on the order of tens of nanometers [18]. Moreover, the intratumoral distribution of both antibodies and nanoparticles is usually highly heterogeneous, with preferential accumulation in perivascular regions [14, 19]. Under such conditions, cells located in close proximity to blood vessels may experience excessive damage, resulting in a local overkill effect, whereas cell populations distant from vascular structures can largely escape exposure to the agent [20].

One of the most widely discussed clinical strategies for overcoming this barrier in the context of antibody–drug conjugates is the co-administration of a “cold” carrier, i.e., unconjugated antibodies that compete with the antibody– drug conjugates for binding sites. Such an approach can shift the receptor saturation front deeper into the tissue and improve the spatial coverage of cells exposed to the therapeutic agent [20, 21]. However, this strategy is effective only within a specific range of ratios between the doses of the cold carrier and the conjugated drug, which in clinical settings may depend, among other factors, on the level of target receptor expression, the lethality threshold, and the physicochemical properties of the agent. In the absence of a prior personalized assessment of the optimal ratio based on available clinical data, this strategy carries the risk of excessive antibody administration, receptor oversaturation, and a dramatic reduction in therapeutic efficacy [20, 21].

A fundamental distinction of radiosensitizing nanoparticles compared to antibody–drug conjugates is that nanoparticles can provide proton dose enhancement even without binding to cellular receptors, which has been confirmed, in particular, by a series of experiments conducted by our group using non-targeted nanoparticles [11, 13, 15, 16]. The conjugation of nanoparticles with tumor-specific antibodies may appear conceptually justified, as it can promote longer retention of the agent within tumor tissue. However, the pronounced spatial heterogeneity in the distribution of targeted nanoparticles within solid tumors raises the question of whether such retention would necessarily translate into increased therapeutic efficacy.

These considerations raise several interrelated questions: does receptor targeting improve the antitumor efficacy of radiosensitizing nanoparticles; can co-administration of unconjugated antibodies enhance treatment efficacy by redistributing nanoparticles within the tumor; what antibody dose is sufficient without being excessive; and when should irradiation be delivered, given that the intratumoral nanoparticle distribution continuously evolves after administration [18]? The answers are expected to vary across patients because they depend on tumor-cell biology, capillary architecture, and the transport kinetics of nanoparticles and antibodies.

The present work addresses these questions from the perspective of mathematical modeling. This approach provides a valuable tool for facilitating the optimization of anticancer treatments by representing the tumor, its microenvironment, and the therapeutic control as a unified dynamic system of equations [22]. Mathematical models make it possible to account for individual variability through simulations performed on cohorts of virtual mice or patients, thereby enabling the assessment of the robustness of therapeutic strategies at the population level. Reliable conclusions obtained using mathematical modeling can subsequently be validated in experiments and clinical trials and can ultimately be integrated into clinical decision-making [23].

## 2. Mathematical model

### 2.1. Equations

The mathematical model used in this study, presented in the form of the system of equations (1) and illustrated in Fig. 1, includes eight variables that are functions of the spatial and temporal coordinates *r* and *t*, as well as two spatially non-distributed variables corresponding to the concentrations of nanoparticles and antibodies in the bloodstream. Their injections, together with the delivery of irradiation, act as external control inputs to the otherwise autonomous system of equations.

**Figure 1:**
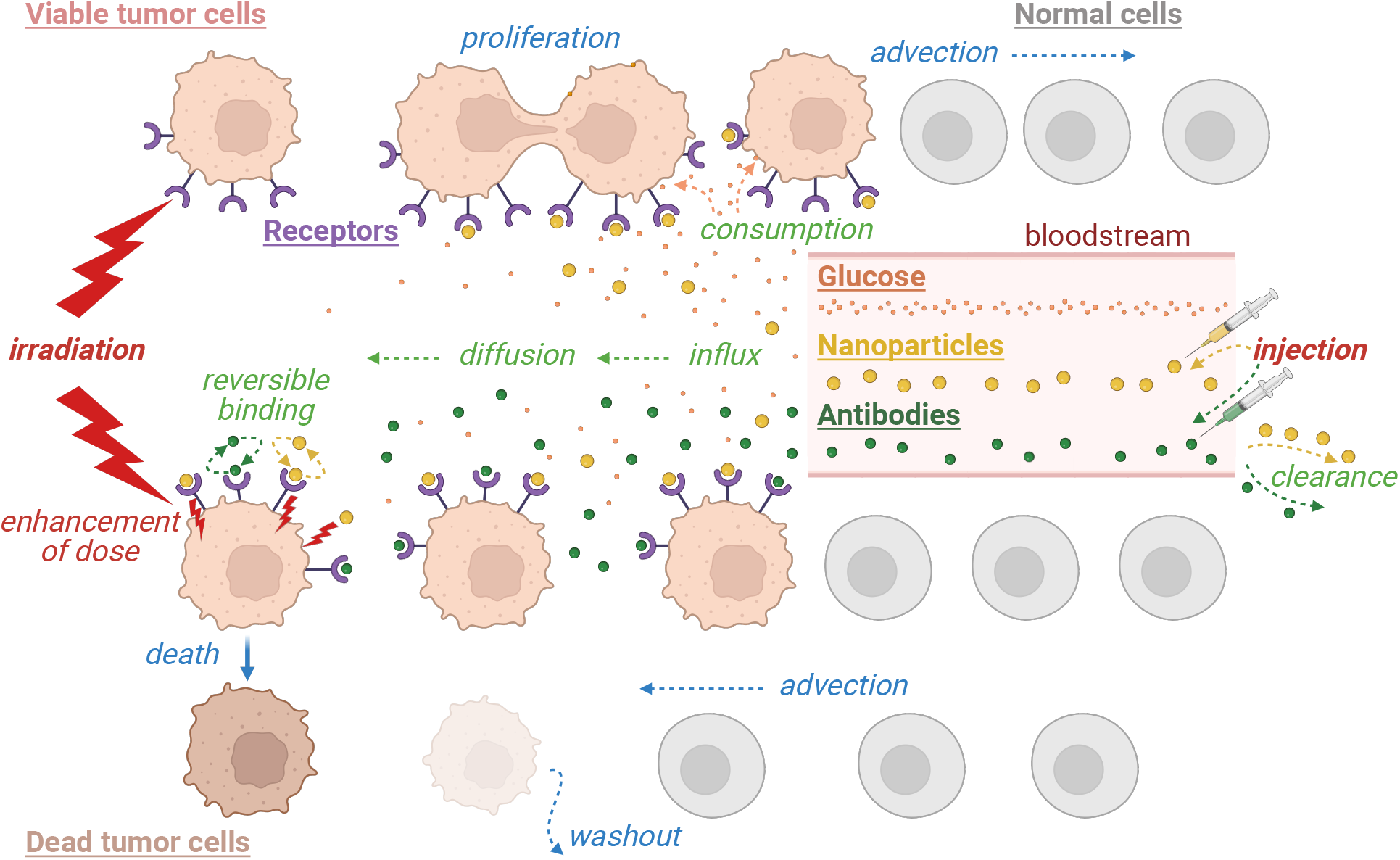
Schematic illustration of the mathematical model described by the system of equations (1). Blue labels correspond to cell dynamics, green labels to substance dynamics, and red labels to therapeutic interventions. Created in BioRender. Kuznetsov, M. (2026) https://BioRender.com/9fec4gt

The model describes spherically symmetric growth of a non-invasive tumor within normal tissue. The proliferation rate of viable tumor cells *n*(*r, t*) depends on the intensity of their glucose consumption, which, in turn, is determined by the local glucose concentration *g*(*r, t*) via a Michaelis–Menten type function. Glucose is explicitly included in the model, as it represents an essential nutrient for the biosynthesis of multiple classes of molecules required for cell division [24]. Tumor cells are surrounded by normal cells *h*(*r, t*). Both tumor and normal tissues are assumed to be fully saturated and incompressible, which is reflected in the condition of constant total cell density normalized to unity. The incompressibility assumption leads to radial cell displacement: as tumor cells proliferate locally, both peripheral tumor cells and surrounding normal cells are displaced outward with velocity *I*(*r, t*), thereby enabling tumor expansion. The modeling of nutrient dynamics is based on our previous work [18]. Glucose enters the tissue from capillaries, whose surface area density is assumed to be proportional to the density of normal tissue. Thus, within the framework of the model, the tumor is assumed to be devoid of its own capillary network. This assumption is based on experimental data showing that functional capillaries are scarce inside the core of large tumors [25]. Glucose diffuses within the tissue and is consumed by viable tumor cells. In the absence of irradiation, the equations for *n, h*, and *g* alone describe free tumor growth.

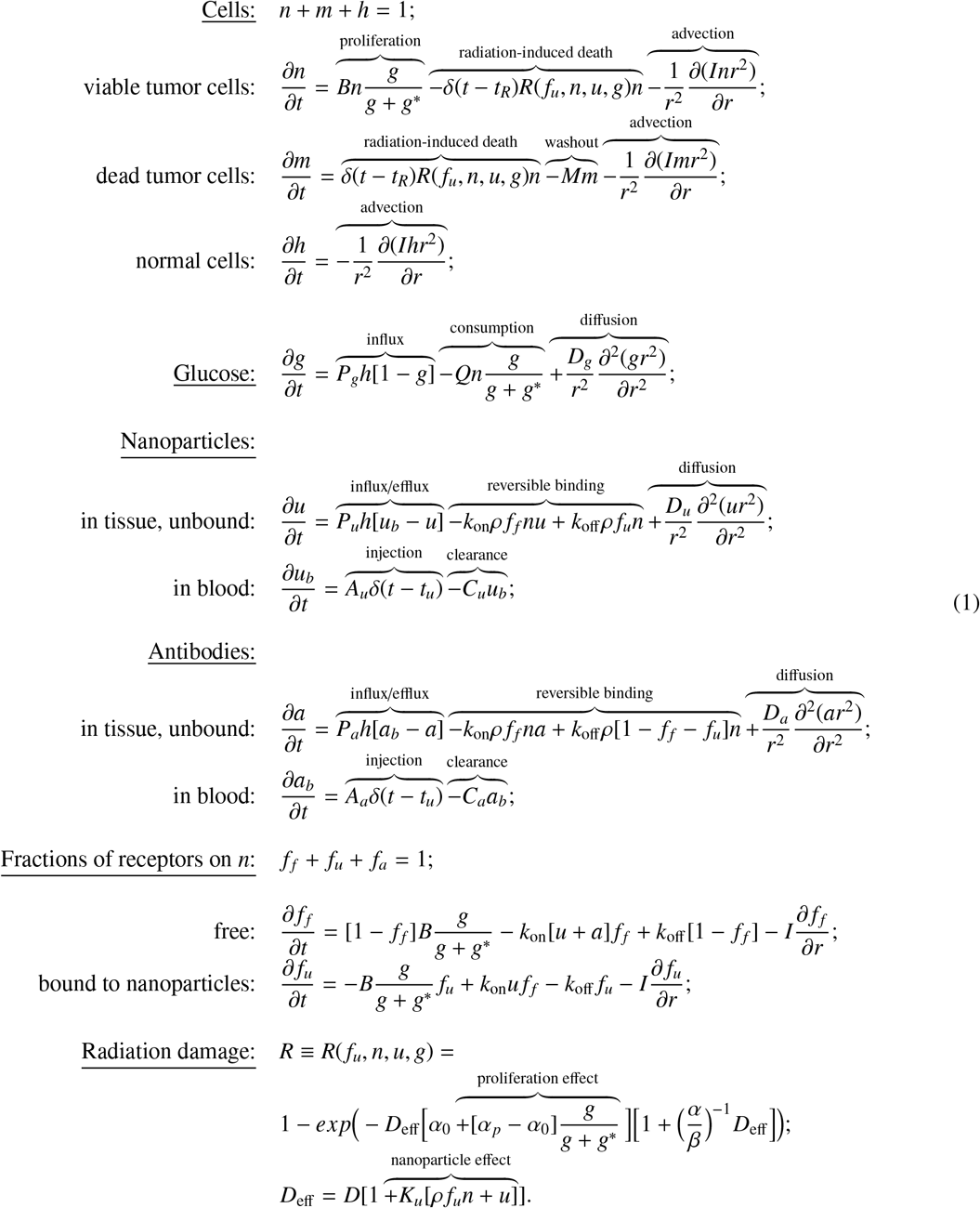

Radiation treatment is modeled as spatially uniform in dose *D*, delivered instantaneously as a single exposure at time *t*_*R*_. However, the radiosensitivity of tumor cells is heterogeneous. In the absence of radiosensitizing nanoparticles, it increases monotonically with the local glucose level, correlating with the metabolic activity of the cells. This assumption is consistent with experimentally observed differences in radiosensitivity between proliferating and quiescent cell populations [26]. In the presence of nanoparticles in the tissue—both bound and unbound to tumor receptors—their local concentration increases the effective radiation dose *D*_eff_. The resulting fraction of tumor cells killed by irradiation, *R*(*r, t*), is modeled using the classical linear–quadratic formalism [27], modified to account for the factors described above. Dead cells *m*(*r, t*) are gradually degraded and washed out from the tumor tissue, affecting the rate of cellular advection and thereby potentially leading to a reduction in tumor volume after irradiation.

Nanoparticles and antibodies are injected into the bloodstream simultaneously at time *t*_*u*_, instantaneously setting their blood concentrations to *u*_*b*_(*t*_*u*_) = *A*_*u*_ and *a*_*b*_(*t*_*u*_) = *A*_*a*_, which subsequently decrease due to systemic clearance. Nanoparticles and antibodies enter the tissue from the bloodstream by diffusion through pores in the capillary walls. The values of diffusive permeability for glucose (*P*_*g*_), antibodies (*P*_*a*_), and nanoparticles (*P*_*u*_) depend on their hydrodynamic radii *p*_*g*_, *p*_*a*_, *p*_*u*_, on their diffusion coefficients and on the spectrum of pores in the capillary walls (see Appendix A). Unbound nanoparticles and antibodies, *u*(*r, t*) and *a*(*r, t*), diffuse within the tissue and can reversibly bind to specific receptors on viable tumor cells. It should be noted that, after prolonged presence within the tissue, free nanoparticles and antibodies can return to the bloodstream through capillaries in normal tissue, as for *u* > *u*_*b*_ and *a* > *a*_*b*_ the corresponding influx terms become negative.

The concentration of specific receptors on tumor cells, ρ, is assumed to be spatially uniform. Receptors can exist in three distinct states. To describe their dynamics, we introduce variables representing the local relative fractions of receptors in each state: free receptors *f* _*f*_ (*r, t*), receptors occupied by nanoparticles *f*_*u*_(*r, t*), and receptors occupied by antibodies *f*_*a*_(*r, t*). Receptors occupied by antibodies are not explicitly tracked in the model due to the constraint that the sum of receptor fractions is equal to unity. During binding, nanoparticles and antibodies compete for the same pool of receptors. Free receptors are expressed on daughter cells during cell division, and the spatial distribution of receptor states depends on the advective velocity *I*, since receptors move in space together with the cellular mass. The derivation of the equations for receptor fractions follows the approach described in our previous work [28] and is presented in Appendix B.

In this study, only a single irradiation event is modeled, after which the variables associated with nanoparticles, antibodies, and receptors no longer influence the equations governing tumor dynamics. We therefore neglect receptors on dead cells and do not explicitly model radiation-induced depletion or transformation of the radiosensitizer, while both effects would need to be considered in extensions to multifraction irradiation protocols.

### 2.2. Numerical methods

The computational code of the model was implemented in C++. The complete source code, processed simulation data underlying the figures and population-level analyses, and the Wolfram Mathematica notebook used for data analysis and plotting are available in the associated research dataset [29]. The glucose equation was treated under a quasi-stationary approximation, justified by the short characteristic timescale of glucose dynamics, and solved at each time step using the Thomas algorithm for tridiagonal systems. For all other variables, the kinetic, diffusion, and advection terms were solved sequentially at each time step. The spatial grid spacing was 5 µm, and the time step was 3.6 s. The kinetic equations were integrated using the explicit Euler method, which is appropriate due to the sufficiently small time step employed. The diffusion terms for nanoparticles and antibodies were solved using the implicit Crank–Nicolson scheme. These classical numerical methods are described in detail, for example, in [30].

The advection equations for cells were solved in conservative form using the method previously proposed by us in [18]. This method implements conservative advection by explicitly redistributing mass between neighboring grid elements and introduces two moving boundary grid points, which allows for sharp tumor–normal tissue interface and outer tissue boundary. This approach is justified in the absence of intrinsic motility of tumor cells and makes it possible to mitigate the effects of artificial diffusion at tissue boundaries that typically arise in classical numerical methods for solving advection equations. Such a formulation ensures the possibility of accurate measurement of the tumor radius *R*_*T*_ (*t*) and the radius of normal tissue *R*_*N*_ (*t*) throughout the simulation. The advection terms for receptor fractions were implemented numerically in an analogous manner, through conservative transport of the local concentrations of receptors in different states, *n f* _*f*_ and *n f*_*u*_, following the same mass redistribution rules as those applied to the tumor cell density *n*.

This approach, applied to the saturated and incompressible tumor and normal tissue in our model (*n* + *m* = 1 ∀*r* ≤ *R*_*T*_ (*t*), *h* = 1 ∀*R*_*T*_ (*t*) < *r* ≤ *R*_*N*_ (*t*)) makes it possible to replace the explicit solution of the equation for normal cells *h* with a substantially less computationally expensive determination of the current radius of normal tissue based on volume conservation as the tumor boundary shifts. It also allows us to avoid explicit computation of the advective velocity *I*(*r, t*) for cells, receptors, and receptor-bound nanoparticles. Nevertheless, this velocity can be calculated, if required, from the following equation, which is obtained by summing the equations for all cell populations in (1) and setting the advective flux to zero at *r* = 0:

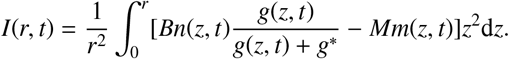

As initial conditions, a spherical region of normal tissue with an initial radius of *R*_*N*_ (0) = 2 cm was considered, with a small spherical colony of tumor cells of radius *R*_*T*_ (0) = 0.2 mm located at its center. The initial glucose concentration profile was specified as its quasi-stationary spatial distribution *g*_*st*_(*r*, 0). The resulting initial conditions were as follows:

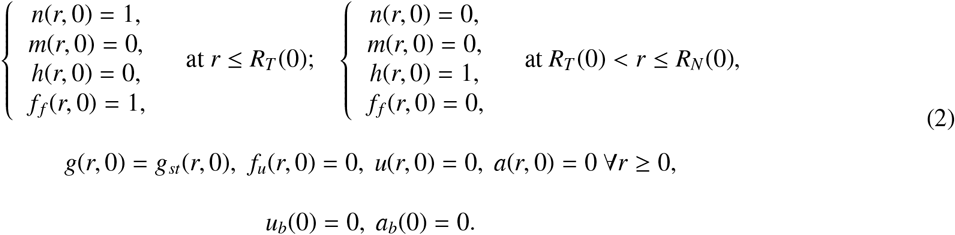

For all spatially distributed variables, zero-flux boundary conditions were imposed:

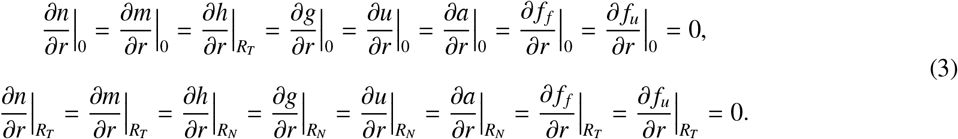

### 2.3. Parameters

The set of model parameters is provided in Table 1. The following normalization parameters were used to obtain their dimensionless model values: 1 h for time; 10^−2^ cm for length; 3 · 10^8^ cells/mL for cell density [31]; 1 mg/mL for glucose concentration (corresponding to the normal glucose concentration in blood [32]); 1 nm for molecular sizes and capillary pore dimensions; 100 cm^2^/cm^3^ for capillary surface density (corresponding to its averaged value for human muscle [33]); 1 Gy for radiation dose; 1 nM = 1 pmol/mL for concentrations of receptors, nanoparticles, and antibodies.

**Table 1:** Model parameters.

| Fixed parameters |  | Value | Normalization | Based on |
| --- | --- | --- | --- | --- |
| $p_g$ | hydrodynamic radius of glucose | 0.36 nm | 0.36 | [34] |
| $p_u$ | hydrodynamic radius of nanoparticles (NPs) | 10 nm | 10 | see text |
| $p_a$ | hydrodynamic radius of antibodies (Abs) | 5 nm | 5 | |
| $C_a$ | Ab clearance rate | 0.14 h<sup>-1</sup> | 0.14 | [35] |
| $D_g$ | glucose diffusion coefficient in tissue | $2.6 \cdot 10^{-6} \frac{\text{cm}^2}{\text{s}}$ | 100 | [36] |
| $D_u$ | NP diffusion coefficient in tissue | $1.6 \cdot 10^{-7} \frac{\text{cm}^2}{\text{s}}$ | 6.5 | see text |
| $D_a$ | Ab diffusion coefficient in tissue | $3.3 \cdot 10^{-7} \frac{\text{cm}^2}{\text{s}}$ | 13 | |
| $g^*$ | Michaelis constant for glucose consumption | 0.04 mM | 0.01 | [37] |
| $\alpha/\beta$ | ratio of radiosensitivity parameters | 10 | 10 | [38] |
| $k_{\text{off}}$ | dissociation rate of NPs and Abs with receptors | $2.2 \cdot 10^{-6} \text{ s}^{-1}$ | 0.08 | see text |
| $K_u$ | dose enhancement coefficient by radiosensitizer | 0.14 nM<sup>-1</sup> | 0.14 | [14] |
| Varied individual parameters |  | Range of variation | Normalization | Based on |
| Variation 1) with respect to capillary pore sizes | | Basic value $\mu = 3$ | | |
| $\mu$ | parameter of capillary pore distribution | 2-4 | 2-4 | Appendix A |
| $P_g = P(D_g, p_g, \mu)$ | capillary permeability for glucose | $2.7 - 10.4 \cdot 10^{-5} \frac{\text{cm}}{\text{s}}$ | 9.8 - 37.7 | |
| $P_u = P(D_u, p_u, \mu)$ | capillary permeability for NPs | $0.06 - 4.1 \cdot 10^{-6} \frac{\text{cm}}{\text{s}}$ | 0.02 - 1.5 | |
| $P_a = P(D_a, p_a, \mu)$ | capillary permeability for Abs | $0.06 - 1.0 \cdot 10^{-5} \frac{\text{cm}}{\text{s}}$ | 0.2 - 3.8 | |
| Variation 2) with respect to metabolic activity of tumor cells | | Basic value $B = 0.03$ | | |
| $B$ | maximum cell proliferation rate | 0.01-0.06 h<sup>-1</sup> | 0.01-0.06 | [31] |
| $Q = 1200 \cdot B$ | glucose consumption rate | $3.2 - 19.2 \cdot 10^{-17} \frac{\text{mol}}{\text{cell} \cdot \text{s}}$ | 12-72 | |
| $M = B$ | washout rate of dead cells | 0.01-0.06 h<sup>-1</sup> | 0.01-0.06 | see text |
| Variation 3) with respect to cell radiosensitivity (RS) | | Basic value $\alpha_0 = 0.05$ | | |
| $\alpha_0$ | linear RS parameter for non-proliferating cells | 0.01-0.1 Gy<sup>-1</sup> | 0.01-0.1 | see text |
| $\alpha_p = 5\alpha_0$ | linear RS parameter for proliferating cells | 0.05-0.5 Gy<sup>-1</sup> | 0.05-0.5 | |
| Variation 4) with respect to receptor expression level | | Basic value $\rho = 250$ | | |
| $\rho$ | density of specific receptors on tumor cells | $10^5$-$10^6 \frac{\text{receptors}}{\text{cell}}$ | 50-500 | [39] |
| Variation 5) with respect to NP clearance rate | | Basic value $C_u = 3$ | | |
| $C_u$ | NP clearance rate | 1-10 h<sup>-1</sup> | 1-10 | see text |
| Therapeutic control |  | Value | Normalization | Based on |
| $D$ | irradiation dose | 30 Gy (or 0) | 30 (or 0) | [14] |
| $A_u$ | NP blood concentration at injection | 10<sup>4</sup> nM (or 0) | 10<sup>4</sup> (or 0) | see text |
| $A_a$ | Ab blood concentration at injection | 10<sup>3</sup> or 10<sup>4</sup> nM (or 0) | 10<sup>3</sup> or 10<sup>4</sup> (or 0) | |
| $k_{\text{on}}$ | association rate of NPs and Abs with receptors | $2.8 \cdot 10^5 \text{ M}^{-1} \text{ s}^{-1}$ (or 0) | 1 (or 0) | |
| $t_u$ | time of NP and Ab administration | $t_u : R_T(t_u) = 1 \text{ cm}$ | | |
| $t_R$ | time of irradiation | $t_u + \Delta t \text{ days}, \Delta t \geq 0$ | | |

The hydrodynamic radius of nanoparticles *p*_*u*_ was estimated based on our *in vivo* experiments [14], in which spherical gold nanoparticles with a core radius of up to 4 nm were used, whose surfaces were pegylated and conjugated with folic acid specific to the tumor receptor folate receptor alpha (FRα), using bovine serum albumin as an intermediate linker. We therefore estimated that these surface modifications increase the effective hydrodynamic radius by ≈6 nm, while not increasing the particle mass by more than several tens of percent.

To estimate antibody-related parameters, we used data for mirvetuximab soravtansine, a clinically approved FRα-targeting antibody–drug conjugate. Its hydrodynamic radius *p*_*a*_ was estimated based on its molecular weight of ≈150 kDa, which is typical for humanized IgG1 monoclonal antibodies, while the clearance rate *C*_*a*_ was derived from its reported half-life of 4.8 days [35]. Due to the lack of corresponding experimental data, the association rate *k*_on_ for targeted nanoparticles and antibodies to receptors was assumed to be identical. Its value was taken from the typical range for IgG–class antibodies, and the dissociation rate *k*_off_ was adjusted to match the known dissociation constant of mirvetuximab soravtansine with FRα, experimentally estimated as 0.08 nM [40]. The dissociation rate was also assumed to be the same for nanoparticles and antibodies given the absence of relevant experimental data.

The diffusion coefficients of nanoparticles and antibodies, *D*_*u*_ and *D*_*a*_, were estimated similarly to our previous studies, assuming an inverse proportionality between the diffusion coefficient of a nanoscale particle and its hydrodynamic radius [18, 41]. The approach used to estimate the range of variation of capillary permeability parameters for different substances was also adopted from our earlier work [18, 41]. This method is described in Appendix A and its outcome is illustrated in Fig. 2. As shown in Fig. 2a), the relative fraction of large pores in capillaries increases with increasing parameter of capillary pore distribution, µ. Consequently, capillary permeability for all considered substances also increases; however, as demonstrated in Fig. 2b), capillary permeability for glucose always exceeds that for antibodies by 1–2 orders of magnitude and that for nanoparticles by 2–3 orders of magnitude.

**Figure 2:**
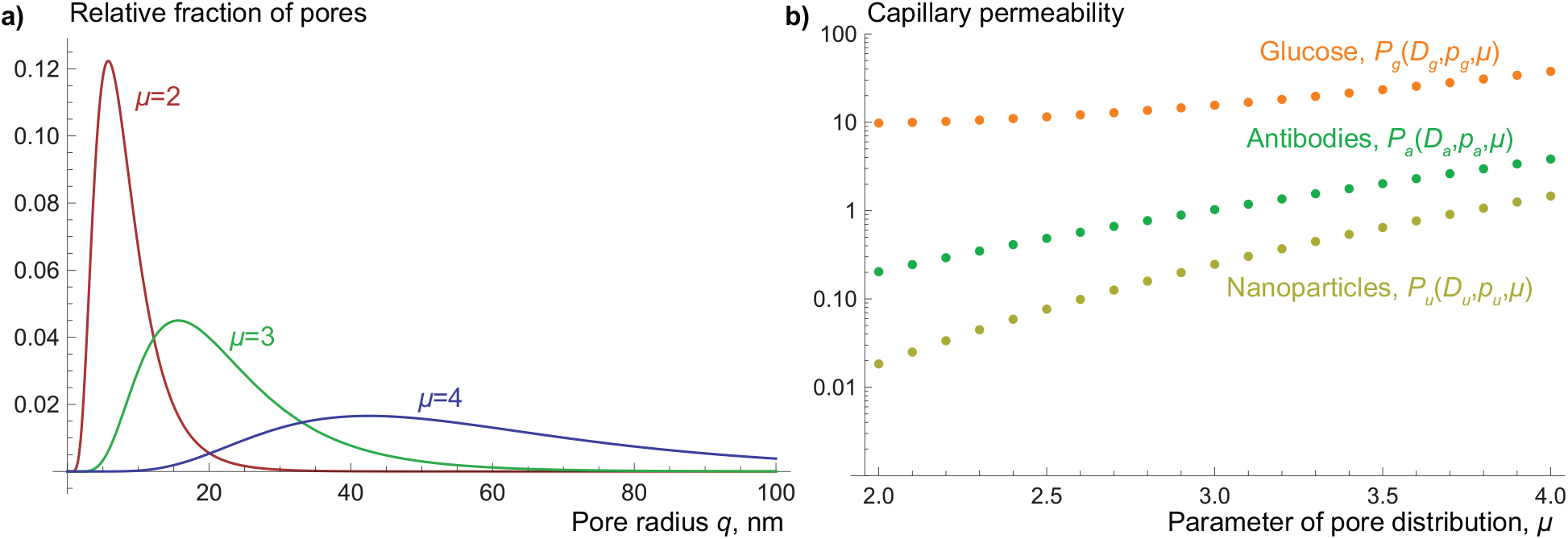
a) The normalized spectra of capillary pore radii used in this study were obtained as lognormal distributions 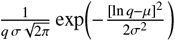, where *q* is the varied pore radius, *σ* = 0.5, and the parameter of capillary pore distribution µ varies from 2 to 4. b) The values of diffusive permeability *P*_*i*_ for glucose (*g*), antibodies (*a*), and nanoparticles (*u*) with set hydrodynamic radii *p*_*i*_ and diffusion coefficients *D*_*i*_ are given by 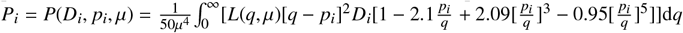 (see Appendix A).

In the parameter variation procedure, the maximum proliferation rate of viable tumor cells *B* was assumed to be proportional to their maximum glucose consumption rate *Q*. The dead-cell washout rate *M* was set equal to the maximum proliferation rate *B*, thereby linking the characteristic timescale of cell loss after irradiation to the timescale of cell-cycle progression [38]. These parameters, as well as the maximum density of tumor cells, were estimated using experimental data from the study [31], performed with radioresistant cells of the murine tumor cell line EMT6/Ro (mammary carcinoma).

The radiosensitive version of the same cell line, EMT6/P, was used in our *in vitro* experiments on proton therapy with gold nanoparticles [14]. Based on these experiments, the reference value of the radiosensitivity parameter for proliferating cells was estimated as α_*p*_ ≈0.1 under the assumption that relatively weak competition for nutrients in a monolayer culture maintains cells in a proliferative state (see the associated research dataset [29]), while in the parameter sweep α_*p*_ was varied over a broader range. The same experimental data were also used to estimate the dose enhancement coefficient of the radiosensitizer *K*_*u*_. The ratio of radiosensitivity parameters α/β was taken to be a typical value for carcinomas [38], while the difference in radiosensitivity between actively proliferating cells and cells experiencing glucose deficiency was chosen based on the fact that the radiosensitivity of quiescent and proliferating cells can differ by a factor of five or more [42].

The range of variation in the concentration of the specific receptor was assessed using published data on the expression of FRα, which is present in a wide variety of epithelial cancers, including breast, lung, kidney, and ovarian tumors [43]. Studies that explicitly reported average numbers of FRα receptors indicate values of 6.9 · 10^5^ receptors per cell (colorectal cancer [44]), 8.3 · 10^5^ receptors per cell (ovarian cancer [45]), and 0.35 pmol of receptors per million cells, corresponding to ≈2.1 · 10^5^ receptors per cell (ovarian cancer [39]).

The characteristic nanoparticle clearance rate *C*_*u*_ was estimated based on our pharmacokinetic data for large nanoparticles [46], including pegylated particles of similar size [47], whose concentration in mouse blood decreases by two orders of magnitude within two hours. However, the effect of albumin coating on the pharmacokinetics of pegylated nanoparticles remains an open question: it may either prolong nanoparticle circulation due to binding to FcRn receptors [48] or shorten it through enhanced opsonization and immune recognition [49]. For this reason, the nanoparticle clearance rate was treated as a varied parameter.

To identify effective nanoparticle-based treatment strategies, we compared six regimens, employing different parameters of treatment control: irradiation dose *D*, nanoparticle and antibody concentrations in blood at the time of injection *A*_*u*_ and *A*_*a*_, and association rate *k*_on_:

1. Free growth (*D* = 0);
2. Radiotherapy (RT) alone (*D* = 30, *A*_*u*_ = 0);
3. RT after injection of non-targeted nanoparticles (*D* = 30, *A*_*u*_ = 10^4^, *k*_on_ = 0);
4. RT after injection of targeted nanoparticles (TNPs) without co-administration of antibodies (Abs) (*D* = 30, *A*_*u*_ = 10^4^, *k*_on_ = 1, *A*_*a*_ = 0);
5. RT after injection of TNPs and moderate dose of Abs (*D* = 30, *A*_*u*_ = 10^4^, *k*_on_ = 1, *A*_*a*_ = 10^3^);
6. RT after injection of TNPs and high dose of Abs (*D* = 30, *A*_*u*_ = 10^4^, *k*_on_ = 1, *A*_*a*_ = 10^4^).

We modeled the injection of a fixed number of nanoparticles into the bloodstream at the moment *t*_*u*_ when tumors reached the same radius of *R*_*T*_ = 1 cm, at which point tumors contain more than one billion cells and pronounced heterogeneity in cell proliferation and radiosensitivity is observed for any admissible set of model parameters.

In our *in vivo* experiments [14], the administered nanoparticle dose was 20 mg, which, assuming a mouse plasma volume of 1 mL and accounting for a gold core radius of 4 nm at a gold density of 19.3 g/cm^3^, corresponds to maximum blood concentration of nanoparticles ≲6000 nM. In the simulations, given the large size of modeled tumors, a higher nanoparticle concentration in the blood at injection was used. No predefined upper bound was imposed on the interval between nanoparticle administration and irradiation when searching for the optimal irradiation time. When modeling antibody administration, the injections of antibodies and nanoparticles were assumed to be simultaneous. The reference value for the antibody concentration in blood upon injection, *A*_*a*_, was based on the standard clinical dose of mirvetuximab soravtansine of 6 mg/kg, which results in a peak blood concentration in adults of ≈900 nM. The single-fraction irradiation dose *D* was chosen to be comparable to that used in our *in vivo* study [14].

## 3. Results

### 3.1. Comparison of regimens for basic values of individual parameters

#### 3.1.1. Free growth and radiotherapy alone

The state of a tumor that has reached a radius of *R*_*T*_ = 1 cm is shown in Fig. 3a. The tumor consists entirely of viable cells; however, only a thin outer layer of these cells has access to glucose levels sufficient to support their active proliferation, as glucose enters the tumor from the surrounding normal tissue via diffusion. The resulting increase in radiosensitivity within this peripheral layer produces a spatially nonuniform distribution of cells surviving irradiation. The overall fraction of surviving cells is ≈0.22%, and in the tumor core the local surviving fraction is slightly higher than this value, whereas at the tumor boundary it is negligibly small. Consequently, after irradiation the outer 0.2 mm layer of the tumor contains no viable cells, while ≈2.8 million surviving cells remain distributed in the deeper tumor regions.

**Figure 3:**
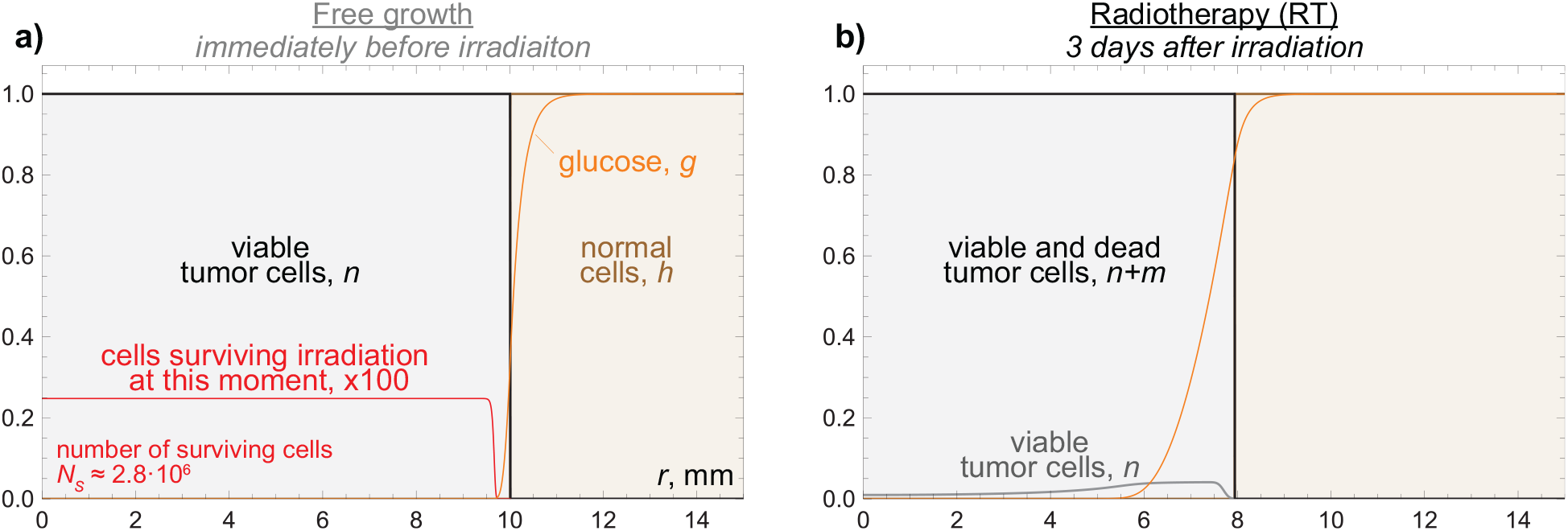
Distributions of model variables obtained by numerical simulation of equations (1) with initial conditions (2) and boundary conditions (3) for the basic parameter values from Table 1, without the use of nanoparticles and antibodies (*A*_*u*_ = 0, *A*_*a*_ = 0). a) Unperturbed tumor growth immediately prior to irradiation. b) Tumor recurrence 3 days after irradiation.

Irradiation eliminates the majority of tumor cells and causes substantial tumor shrinkage, as illustrated in Fig. 3b, corresponding to the third day after irradiation. During this time the tumor radius decreases by ≈2 mm. However, the reduction of competition for glucose among the small number of remaining viable tumor cells results in a substantial increase in glucose availability within the tumor. Therefore, in the first days after irradiation most of the surviving tumor cells proliferate at nearly maximum rate *B*. As a result, by the third day after therapy their number increases almost sevenfold, and the tumor begins to regain its original structure, with glucose levels again falling to low values in the tumor core and proliferating cells becoming localized in the outer tumor layer. By day 11 after irradiation the tumor radius starts to increase, and it returns to its pre-irradiation value of *R*_*T*_ = 1 cm by day 42.

#### 3.1.2. Administration of nanoparticles without antibodies

Figure 4 compares tumor dynamics and the nanoparticle penetration under the basic parameter set for the therapy regimens involving injection of nanoparticles at moment *t*_*u*_, when tumor radius reaches 1 cm (see Table 1). Administration of nanoparticles in our model results in their maximal blood concentration of *u*_*b*_(*t*_*u*_) = *A*_*u*_ = 10^4^ nM. If this level could be maintained in the blood for a prolonged period, then for the chosen capillary permeability of nanoparticles of *P*_*u*_ ≈0.25 the nanoparticle concentration in normal tissue *u*(*R*_*N*_) would reach *A*_*u*_/2 within ≈3 hours. However, *u*(*R*_*N*_) does not reach such high values due to the very rapid clearance of nanoparticles from the blood, whose half-life is less than 15 minutes. Already within the first hour after injection, *u*(*R*_*N*_) and *u*_*b*_ become comparable at the level of several hundred nM, and subsequently the nanoparticle concentration in normal tissue decreases as a result of their net efflux back into the bloodstream. Nevertheless, nanoparticles entering tumor tissue by diffusion are retained there for a substantial period, being unable to return directly to the bloodstream due to the absence of tumor capillaries.

**Figure 4:**
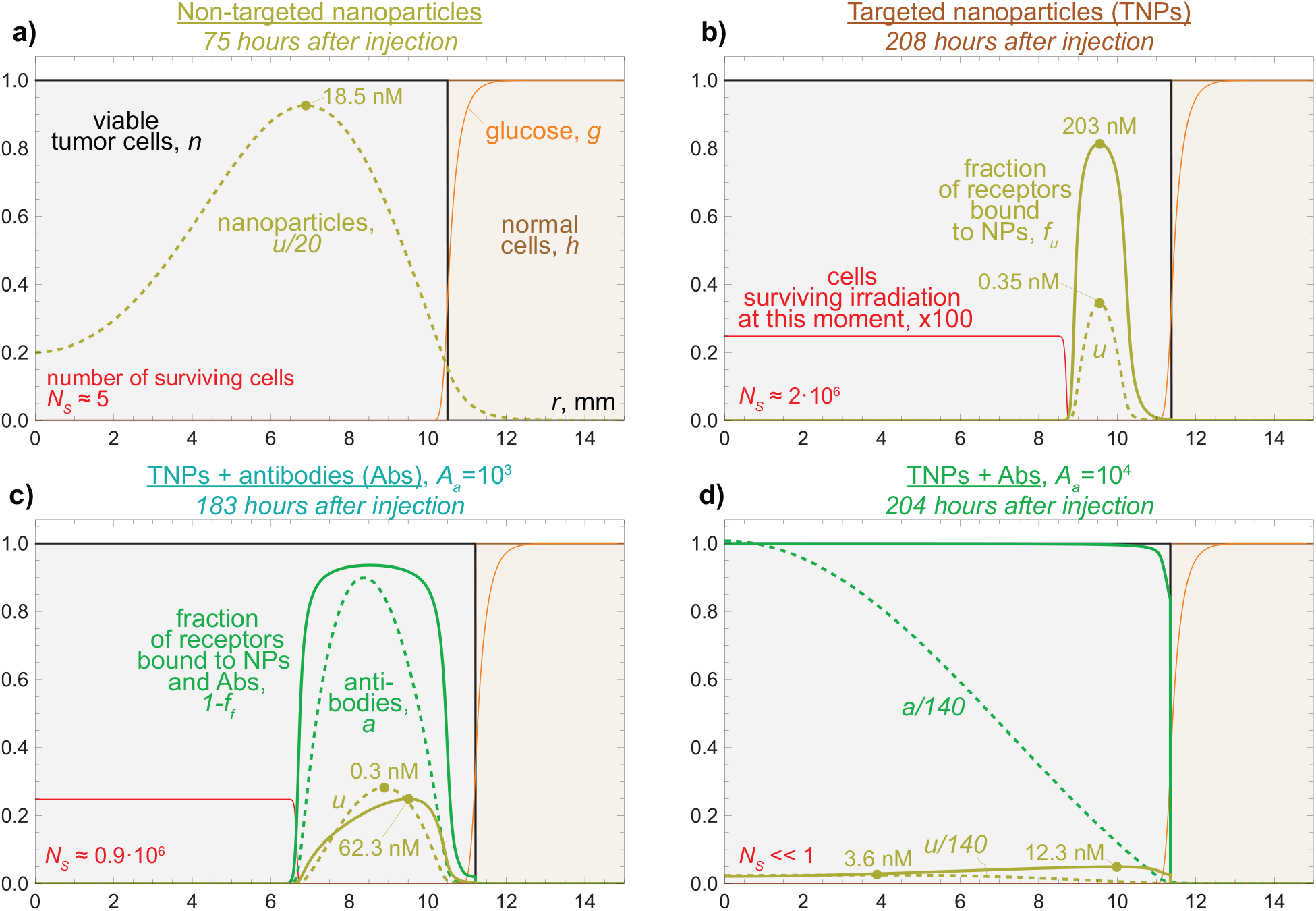
Distributions of model variables for the basic parameter values from Table 1 and for different therapy regimens specified by the nanoparticle concentration in blood at the time of injection *A*_*u*_, the rate of association of nanoparticles and antibodies with tumor receptors *k*_on_, and the antibody concentration in blood at the time of injection *A*_*a*_. The figures illustrate nanoparticle penetration at the time points when irradiation results in minimization of surviving tumor cells for: a) non-targeted nanoparticles that do not bind to tumor receptors (*A*_*u*_ = 10^4^, *k*_on_ = 0, *A*_*a*_ = 0); b) targeted nanoparticles binding to tumor receptors (*A*_*u*_ = 10^4^, *k*_on_ = 1, *A*_*a*_ = 0); c) targeted nanoparticles with simultaneously administered antibodies binding to the same receptors at a moderate antibody dose (*A*_*u*_ = 10^4^, *k*_on_ = 1, *A*_*a*_ = 10^3^); d) targeted nanoparticles with simultaneously administered antibodies at high dose (*A*_*u*_ = 10^4^, *k*_on_ = 1, *A*_*a*_ = 10^4^).

Thus, the “enhanced permeability and retention” effect characteristic of solid tumors [50] ensures the maintenance of a substantial level of nanoparticles in the tumor and the adjacent tissue for several days, even in the absence of nanoparticle binding to tumor receptors, as demonstrated in Fig. 4a. As can be seen from this figure, 75 hours after administration of non-targeted nanoparticles their concentration in the tumor lies in the range of ≈3–18.5 nM, providing effective cell killing throughout the entire tumor volume. However, such therapy leaves ≈5 tumor cells viable, which ultimately leads to regrowth of the tumor radius to 1 cm within 74 days after nanoparticle injection.

The presence of binding of nanoparticles to tumor receptors enhances their retention within the tumor, but leads to a less effective irradiation outcome. As shown in Fig. 4b, corresponding to 208 hours after nanoparticle administration, binding of nanoparticles substantially restricts their penetration into the tumor core. As a result, in the central tumor zone with a radius of ≈8.5 mm radiosensitization is negligible. Moreover, due to the continued proliferation of tumor cells in the outer tumor layer following nanoparticle administration, this region also becomes largely depleted of nanoparticles by this time point; however, effective killing of these actively proliferating cells is ensured by their increased intrinsic radiosensitivity. The illustrated time point minimizes the total number of surviving tumor cells. At later times, as cells continue to proliferate, in regions with low nanoparticle and glucose concentrations (corresponding to distance from tumor center *r* ≈11–11.5 mm), a second zone of tumor cells with comparatively low radiosensitivity emerges. When irradiation is performed at the chosen time point, despite a markedly higher maximal local concentration of targeted nanoparticles in the tumor (≈200 nM) compared to the case with non-targeted nanoparticles (≈18.5 nM, Fig. 4a), the use of targeted nanoparticles in this model simulation reduces the surviving fraction of tumor cells only by approximately a factor of two relative to irradiation alone (Fig. 3a), due to the limited spatial coverage of tumor cells by nanoparticles. This therapy regimen results in a delay of tumor regrowth by ≈57 days.

#### 3.1.3. Co-administration of nanoparticles and antibodies

Co-administration of a moderate dose of antibodies together with nanoparticles expands the spatial extent of nanoparticle receptor occupancy, as demonstrated in Fig. 4c, which corresponds to the optimal irradiation time point at 183 hours after simultaneous administration of nanoparticles and antibodies, with the maximal antibody concentration in blood being an order of magnitude lower than that of nanoparticles (*A*_*a*_ = *A*_*u*_/10 = 10^3^). Despite their tenfold lower initial blood concentration, the slower clearance, higher capillary permeability, and faster diffusion of antibodies result in ≈3.6 times more receptor-bound antibodies than receptor-bound nanoparticles at this time point. Competition for receptors between antibodies and nanoparticles also leads to reduced nanoparticle retention within the tumor: compared to the therapy regimen without antibodies (Fig. 4b), the total number of bound nanoparticles in the tumor in this case is lower by a factor of ≈1.7. However, receptor competition also increases the penetration depth of nanoparticles, which under this regimen are able to diffuse farther from the tumor boundary before becoming bound to receptors. This effect plays a decisive role in improving irradiation efficacy, reducing the radius of the tumor core where the radiosensitization effect is virtually absent by ≈2 mm compared to the case without antibodies. This therapy regimen results in a delay of tumor regrowth by ≈59 days.

Increasing the antibody dose to a level equal to that of nanoparticles (*A*_*a*_ = *A*_*u*_ = 10^4^) allows the benefit of nanoparticle redistribution for therapeutic effect to be substantially enhanced, as shown in Fig. 4d, corresponding to the optimal irradiation time point at 204 hours after administration of nanoparticles and antibodies. Owing to the high antibody dose, tumor receptors become nearly saturated throughout the entire tumor volume, with the exception of the outer tumor layer, where, due to ongoing cell proliferation, the local fraction of unbound receptors reaches ≈16%. The concentration of bound nanoparticles within the tumor volume in this case varies from ≈5.5 to ≈12.3 nM. An additional contribution to the increased irradiation efficacy is provided by unbound nanoparticles that remain in the tumor due to the enhanced permeability and retention effect. As a result, the total nanoparticle concentration within the tumor volume lies in the range of ≈7–13 nM (corresponding to a gold concentration of ≳20 µg/mL). Sustained enhancement of radiosensitization is maintained through reversible binding and gradual diffusive efflux of free antibodies from the tumor, which leads to progressively more nanoparticles (having a lower diffusion coefficient) becoming bound to tumor receptors. Irradiation at this time point reduces the predicted number of surviving cells below 10^−4^, well below the deterministic cure threshold used in this study.

#### 3.1.4. System dynamics in different regimens

The model dynamics for the simulations described above are demonstrated in Fig. 5. The temporal evolution of the potential irradiation efficacy after nanoparticle administration is illustrated in Fig. 5a through plots of the number of tumor cells *N*_*s*_ that would survive if irradiation was performed at the current time point.

**Figure 5:**
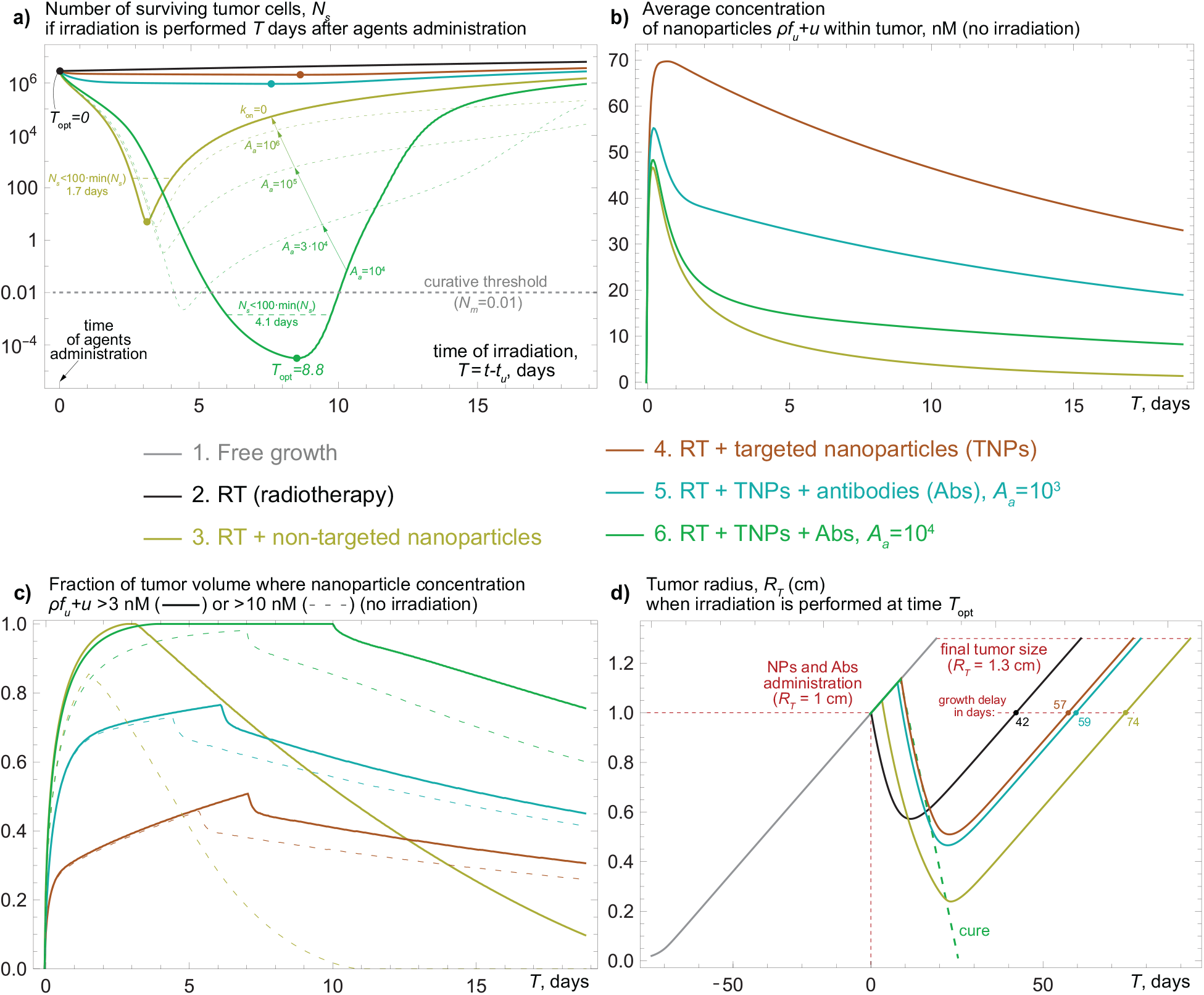
Dynamics of the model simulations illustrated in Figs. 3 and 4. a) Number of tumor cells that would survive, *N*_*s*_, if irradiation was performed at the current time point (in days) after nanoparticles and antibodies are administered, *T* = *t* − *t*_*u*_. Dots mark the optimal time points for irradiation, *T*_opt_, corresponding to the minimal number of surviving cells, min(*N*_*s*_). Dashed lines visualize the fact that with the increase of concentration of injected antibodies, *A*_*a*_, the system effectively approaches the regime of non-targeted nanoparticles (*k*_on_ = 0). b) Temporal evolution of the average nanoparticle concentration within the tumor volume in regimens involving nanoparticle administration. c) Fraction of tumor volume in which the total concentration of bound and unbound nanoparticles exceeds the selected thresholds (3 nM and 10 nM) for the same regimens. d) Tumor radius, *R*_*T*_, for different regimens with irradiation performed at the corresponding optimal time points *T*_opt_.

For irradiation alone, in the absence of nanoparticles, the most effective option in terms of minimizing the number of viable tumor cells is to perform therapy as early as possible, since the efficacy of delayed irradiation is compromised by the increase in tumor cell number during the waiting period (however, this option does not guarantee maximization of the tumor regrowth delay and, from that perspective, should be less advantageous than later irradiation, as demonstrated by numerous modeling studies, e.g., [51]). When nanoparticles are administered, the curves exhibit pronounced minima corresponding to time points when nanoparticle accumulation—providing radiosensitization of both existing and newly formed tumor cells—creates optimal conditions for tumor irradiation.

The primary determinant of irradiation efficacy is the spatial coverage of tumor tissue by nanoparticles rather than their maximum or average intratumoral concentration. As shown in Fig. 5b, in every considered regimen involving nanoparticle administration, their average concentration in tumor tissue reaches its peak already within the first day after injection. However, minimization of the number of viable tumor cells is achieved substantially later in every case (Fig. 5a). Moreover, tumor cell kill does not correlate with the average nanoparticle concentration: while the co-administration of antibodies reduces the total amount of nanoparticles penetrating into the tumor, it nevertheless results in a pronounced decrease in *N*_*s*_.

Instead, irradiation efficacy benefits from sufficient spatial coverage of tumor cells by nanoparticles. This relationship is illustrated in Fig. 5c. Comparison of panels a and c demonstrates that irradiation is most effective when a large fraction of the tumor is exposed to sufficiently high local nanoparticle concentrations. For targeted nanoparticles without antibodies, the minimal number of surviving cells, min(*N*_*s*_), is reached near day 9, when less than half of the tumor volume exhibits nanoparticle concentrations above 3 nM. In this case, min(*N*_*s*_) decreases by only ≈30% compared to irradiation alone. Co-administration of antibodies shifts the optimal irradiation time only moderately, but markedly improves both spatial coverage of tumor cells by nanoparticles and irradiation efficacy. At a moderate antibody dose of *A*_*a*_ = 10^3^, when approximately two thirds of tumor volume has nanoparticle concentration above 3 nM, min(*N*_*s*_) decreases by ≈70% compared to irradiation alone. A higher antibody dose, *A*_*a*_ = 10^4^, results in nearly complete tumor coverage with nanoparticle concentration above 3 nM and leads to a dramatic reduction in the number of surviving tumor cells.

The use of non-targeted nanoparticles also improves spatial coverage of cells by nanoparticles and treatment efficacy, but shifts the optimal irradiation time to the beginning of day 4, which is due to the subsequent rapid unconstrained diffusive efflux of unbound nanoparticles from the tumor. Notably, compared with non-targeted nanoparticles, targeted nanoparticles combined with a high antibody dose not only achieve greater tumor cell kill but also create a prolonged window of enhanced therapeutic efficacy, associated with increased nanoparticle retention within the tumor.

Importantly, Fig. 5a illustrates the conceptual advantage of co-administration of antibodies with radiosensitizing nanoparticles. As the administered antibody dose increases beyond *A*_*a*_ = 10^4^, receptor binding of nanoparticles is progressively suppressed, and in the limit of very high antibody concentrations the system effectively approaches the regime of non-targeted nanoparticles (*k*_on_ = 0). Therefore, in this setting, excessive antibody dosing cannot reduce tumor cell kill below the level achievable with non-targeted nanoparticles, which still yields greater tumor cell kill than the use of targeted nanoparticles without antibodies.

The superiority of non-targeted over targeted nanoparticles in maximizing tumor cell kill is not universal across arbitrary parameter sets. Nevertheless, as demonstrated in Appendix C, it is expected to manifest itself in single-high-dose settings. In contrast, the use of targeted nanoparticles without co-administration of antibodies may outperform non-targeted nanoparticles in terms of tumor cell kill in the regime of sufficiently low irradiation dose (or sufficiently high intrinsic radioresistance of tumor cells) and low nanoparticle dose. Such behavior may therefore become more relevant for low-dose fractions characteristic of fractionated radiotherapy, which is not considered in the present study, but this qualitative conclusion is conceptually consistent with our previous results on the spatial optimization of fractionated irradiation, where low-dose fractions of irradiation were shown to benefit from preferential dose localization near the tumor rim [52].

The tumor growth curves for the considered cases with irradiation performed at the time point corresponding to the minimal number of surviving cells are presented in Fig. 5d. As can be seen from this figure, the delay in tumor regrowth in this setting correlates with the reduction in the number of tumor cells surviving after irradiation. In the present study we employ a continuous modeling framework, in which the number of cells is represented as a real-valued quantity, which leaves some flexibility in the interpretation of tumor cure scenarios that are traditionally defined using probabilistic approaches [53]. To retain a deterministic framework, we classified a tumor as cured when the predicted number of viable tumor cells fell below 0.01. Under a classical Poisson interpretation, this threshold corresponds to a tumor cure probability of *e*^−0.01^ ≈0.99 [54]. Under this definition, the scenario involving the use of a high antibody dose, *A*_*a*_ = 10^4^, results in tumor cure within the modeling framework.

### 3.2. Comparison of treatment regimens under parameter sweep

To assess the efficacy of different treatment regimens across a broad range of physiologically plausible parameters, we performed simulations for 100 “virtual mice,” each characterized by a distinct set of model parameters. Specifically, the maximum tumor cell proliferation rate *B*, the parameter of the capillary pore size distribution µ, the baseline linear radiosensitivity α_0_, the surface density of specific receptors ρ, and the nanoparticle clearance rate *C*_*u*_ were sampled independently from uniform distributions over the ranges listed in Table 1. The complete set of sampled parameter combinations is provided in the associated research dataset [29].

As previously, irradiation efficacy was first quantified by the minimal number of surviving tumor cells, min(*N*_*s*_), achieved when irradiation was applied at the personalized optimal time points 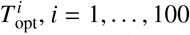, after nanoparticle and antibody administration (see Fig. 5a). The regimen involving targeted nanoparticles administered together with a high antibody dose (*A*_*u*_ = 10^4^, *k*_on_ = 1, *A*_*a*_ = 10^4^) remained the most effective in terms of potential tumor cell kill for 90% of the virtual mice, providing a window for curative irradiation in 15 cases. Moreover, this regimen yielded the smallest value of min(*N*_*s*_) for each of the 68 mice whose receptor density lay in the range ρ ∈ [80, 400], corresponding to ≈1.6 − 8 · 10^5^ receptors per cell. For six mice with receptor densities below this interval, the regimen with targeted nanoparticles and a lower antibody dose (*A*_*u*_ = 10^4^, *k*_on_ = 1, *A*_*a*_ = 10^3^) produced the smallest min(*N*_*s*_). Conversely, for four mice with receptor densities above this range, the optimal strategy was the use of non-targeted nanoparticles (*A*_*u*_ = 10^4^, *k*_on_ = 0), which, as discussed above, can be regarded as the limiting case of *k*_on_ = 1 and *A*_*a*_ → ∞. For each of these ten mice, the high-antibody-dose regimen remained the second-best option.

Crucially, the regimen involving targeted nanoparticles without antibody co-administration (*A*_*u*_ = 10^4^, *k*_on_ = 1, *A*_*a*_ = 0) was inferior to all other nanoparticle-based regimens in terms of tumor cell kill for each of the 100 virtual mice. Taken together, these results support the conclusion that, although the optimal antibody dose should depend on individual physiological parameters—with the receptor density ρ being a particularly influential factor—the co-administration of antibodies by itself provides a robust enhancement of tumor cell kill across a physiologically heterogeneous population.

These results, however, correspond to an idealized scenario in which irradiation is administered at the personalized optimal time point, which cannot be known *a priori*. Moreover, they leave open the question of whether a reduction in the number of surviving tumor cells translates robustly into a longer delay of tumor regrowth for non-cured tumors, which may be regarded as an additional measure of treatment efficacy and which optimization would, in general, require a different approach.

To address these issues, we performed an *in silico* trial for the same 100 virtual mice using a single fixed interval between agent administration and irradiation, *T*_*R*_ = *t*_*R*_ − *t*_*u*_, for all mice within a given treatment regimen. For the regimen involving targeted nanoparticles co-administered with a high antibody dose, *A*_*a*_ = 10^4^, the optimal universal interval was found to be *T*_*R*_ = 6 days. This choice resulted in tumor cure in 14 out of the 15 mice that could potentially be cured under this regimen, as illustrated in Fig. 6. Such a relatively long waiting period before irradiation allows sufficient time for spatial redistribution of nanoparticles, thereby improving tumor coverage (see Fig. 4d). By contrast, using *T*_*R*_ = 3 days in this regimen did not yield any tumor cures. For the regimen employing non-targeted nanoparticles, the optimal universal interval was shorter, *T*_*R*_ = 4 days, owing to the reduced retention of nanoparticles within the tumor. With this choice of *T*_*R*_, tumor cure was achieved in 5 out of the 7 mice that could potentially be cured in this regimen.

**Figure 6:**
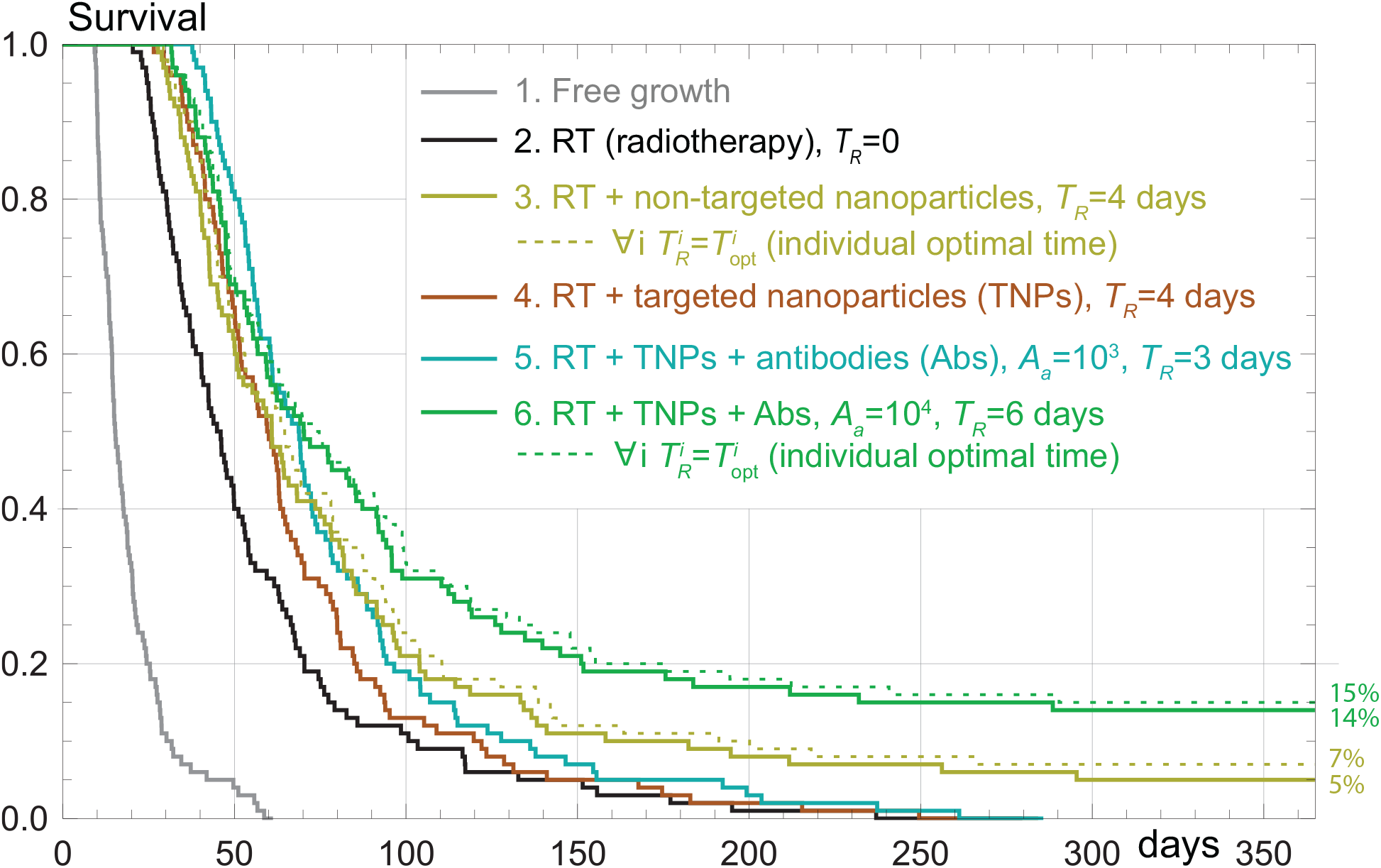
Survival curves from an *in silico* trial of 100 virtual mice, each defined by a randomly sampled set of five individual parameters (Table 1) and simulated under five treatment regimens (solid curves). Nanoparticles and antibodies were administered when the tumor reached a radius of 1 cm. Survival was defined as the fraction of virtual mice whose tumors had not reached the endpoint radius of 1.3 cm within 365 days. For each regimen, a common interval *T*_*R*_ between agent administration and irradiation was restricted to an integer number of days and selected to maximize the number of tumor cures or, when no cures were achieved, the cumulative time to the tumor-size endpoint. Solid curves are compared with best-case personalized benchmarks obtained by irradiating each virtual mouse at its individual optimal time 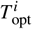, *i* = 1, …, 100 (dashed curves).

Two other regimens, involving administration of targeted nanoparticles without antibodies or with a moderate antibody dose, *A*_*a*_ = 0 and *A*_*a*_ = 10^3^, were found to have no potential to achieve tumor cure, irrespective of the interval between agent administration and irradiation. For these regimens, we therefore selected the universal time intervals *T*_*R*_ that maximized the sum of tumor regrowth times up to the final tumor size *R*_*T*_ = 1.3 cm.

As shown in Fig. 6, despite their inferior long-term performance, these regimens transiently provide comparable or even better survival than the previously discussed regimens during the first two months after treatment initiation. In particular, for nearly half of the mice irradiated on day 4 after nanoparticle injection, administration of targeted nanoparticles without antibodies resulted in a slightly longer time to the tumor-size endpoint than administration of non-targeted nanoparticles. This observation indicates that an increase in tumor cell kill does not necessarily translate into a longer tumor regrowth delay: localized elimination of a substantial fraction of cells near the tumor rim may impair tumor regrowth more effectively than elimination of a larger number of cells distributed throughout the tumor volume, especially in the case of rapidly-growing radioresistant tumors.

Nevertheless, for every virtual mouse, the addition of a moderate antibody dose, *A*_*a*_ = 10^3^, to targeted nanoparticles did lead to an increase in tumor regrowth delay. Overall, these results indicate that antibody co-administration yields a robust improvement in survival across a physiologically heterogeneous population. While moderate antibody doses consistently prolong tumor regrowth time, high antibody doses may slightly shorten the time to the tumor-size endpoint in a subset of virtual mice but produce a pronounced and robust increase in tumor cure probability.

## 4. Discussion

Despite its favorable dose-distribution characteristics relative to conventional photon radiotherapy, proton therapy has shown little or no advantage in many clinical settings [55, 56]. Further methodological advances are therefore required to unlock the full therapeutic potential of this high-cost modality and to justify its broader clinical adoption. One promising approach is the use of radiosensitizing nanoparticles, which, upon accumulation within the tumor, can increase the effective deposited dose through proton-induced interactions, while limiting additional normal-tissue toxicity through preferential tumor accumulation.

The efficacy of radiosensitizing nanoparticles is, however, limited by their restricted transport into tumor tissue through the capillary network—an effect that has been observed experimentally [14] and is intrinsic to nanoscale therapeutic agents in general [20]. This restricted delivery leads to preferential nanoparticle accumulation in perivascular regions and the formation of localized zones of excessive irradiation (resulting in cellular “overkill”), while the tumor interior remains insufficiently sensitized.

In the present study, we demonstrate by means of mathematical modeling that this limitation can be mitigated through the co-administration of targeted nanoparticles and antibodies binding to the same tumor-specific receptors. Following administration, a waiting period of several days allows for progressive spatial redistribution of nanoparticles within the tumor, leading to the emergence of a prolonged window of enhanced therapeutic efficacy. Delivering a single high-dose irradiation within this window is predicted to significantly increase tumor cell kill. This effect arises because the overall efficacy of radiosensitization is governed primarily by the spatial coverage of tumor tissue by a sufficient amount of nanoparticles at the time of irradiation, rather than by the maximum or average nanoparticle concentration achieved within the tumor.

The optimal dose of antibodies is expected to depend on the physicochemical properties of both nanoparticles and antibodies, as well as on individual physiological parameters, and therefore may be difficult to personalize. An important and conceptually non-trivial implication of our results, however, is the fundamentally asymmetric risk associated with antibody co-administration in the context of radiosensitizing nanoparticles. In contrast to antibody–drug conjugates, for which excessive dosing of unconjugated antibodies may cause receptor oversaturation and a dramatic loss of therapeutic efficacy [20, 21], thereby posing a serious obstacle to clinical implementation, the situation for nanoparticle-mediated radiosensitization is qualitatively different in the context of single high-dose irradiation. In the limit of very high antibody doses, the system effectively approaches the regime of non-targeted nanoparticles. In the high-dose setting, this limiting non-targeted regime remains more effective in terms of tumor cell kill than targeted nanoparticles administered alone, owing to the improved spatial coverage of tumor tissue by nanoparticles. This observation indicates that antibody co-administration for radiosensitizing nanoparticles has a built-in safety margin, offering a favorable therapeutic scenario that is expected to provide a robust treatment benefit despite substantial variability in tumor characteristics.

We emphasize, however, that the use of a single high-dose irradiation serves here as a model scenario whose conclusions are intended to be verified in experimental settings and does not aim to reproduce a specific clinical protocol. The investigation of multi-fraction irradiation schemes, including effects associated with prolonged treatment such as immune responses and vascular remodeling, represents a natural direction for future extensions of the model.

Notably, the radiosensitivity and radiosensitization parameters employed in the model were informed by our own experimental measurements [14] as well as published literature data, grounding the simulations in physiologically plausible quantitative ranges. Taken together, the results provide a quantitative framework for the rational design and subsequent experimental testing of nanoparticle-assisted radiotherapy protocols.

## Supporting information

Source code, processed simulation data, and analysis files

## Appendix A. Estimation of capillary permeabilities

The exchange of small lipid-insoluble substances between blood and interstitial fluid occurs primarily via passive diffusion through capillary pores. This process is described by Fick’s law, which for the transport of molecules with hydrodynamic radius *p* through pores modeled as cylindrical channels of radius *q*, oriented perpendicular to the capillary surface, takes the form:

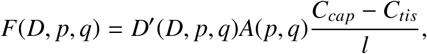

where *F*(*D, p, q*) is the flux of the considered substance, *D*^′^(*D, p, q*) is its diffusion coefficient inside the pore, which depends on the free diffusion coefficient of the substance *D*, (*C*_*cap*_ − *C*_*tis*_) is the difference between the concentrations of the substance in capillary blood and interstitial fluid, *l* is the pore length, and *A*(*p, q*) is the fraction of the pore cross-sectional area accessible for penetration by the substance molecules.

Since a molecule experiences steric exclusion, meaning that it cannot approach the pore wall closer than its hydrodynamic radius, the area accessible to molecules in *N* pores is equal to

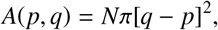

Diffusion of substances through pores is additionally restricted by hydrodynamic resistance, which can be quantitatively described using the empirically derived Renkin equation [57]:

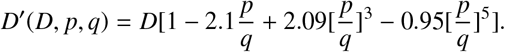

To describe the capillary system, it is convenient to introduce a physically measurable parameter — the permeability *P*(*D, p, q*), defined as follows:

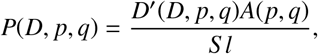

where *S* is the volumetric density of capillary surface area. This definition leads to the following equation for substance transport:

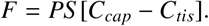

Although real pores are not ideal cylinders, it has been shown that pores in skeletal and cardiac muscle restrict diffusion approximately in the same way as cylindrical pores with a radius of 4–5 nm [58]. Capillaries formed as a result of tumor angiogenesis, as well as those exposed to vascular endothelial growth factor (VEGF) secreted by tumor cells under metabolic stress, contain much larger openings in their walls, known as fenestrations, whose cross-sectional size reaches tens of nanometers [33].

In the present work, we assume that the spectrum of capillary pore radii is described by a lognormal distribution of the form

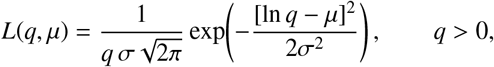

where σ = 0.5, and the parameter µ varies from 2 to 4, which yields the pore radius spectra shown in Fig. 2a).

To account for transport through a distribution of pore sizes, we define the effective permeability as

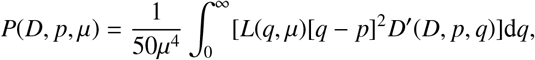

where the coefficient 1/(50µ^4^), which normalizes the total area of all pores, was chosen to obtain physiologically reasonable values of tumor capillary permeability for glucose [41], while ensuring that an increase in µ leads to an increase in the overall capillary permeability for the considered substances. The resulting permeability values for glucose, antibodies, and nanoparticles, illustrated in Fig. 2b), are determined through their hydrodynamic radii and diffusion coefficients as *P*_*g*_ = *P*(*D*_*g*_, *p*_*g*_, µ), *P*_*a*_ = *P*(*D*_*a*_, *p*_*a*_, µ), and *P*_*u*_ = *P*(*D*_*u*_, *p*_*u*_, µ).

## Appendix B. Derivation of the equations for receptor fractions

We consider a single population of viable tumor cells with density *n*(*r, t*). Each cell carries ρ specific receptors. The total number of receptors per unit volume of tumor tissue is therefore ρ*n*(*r, t*).

Receptors can exist in three states:

- free (index *f*),
- bound to nanoparticles (NPs) carrying the radiosensitizer (index *u*),
- bound to “cold” antibodies (Abs) (index *a*).

Let us introduce the absolute concentrations of receptors

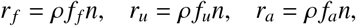

where the variables *f* _*f*_, *f*_*u*_, and *f*_*a*_ denote the fractions of the corresponding receptors on viable tumor cells, such that

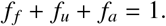

Following the assumptions of the model, the equations for the concentrations of receptors on viable tumor cells take the following form:

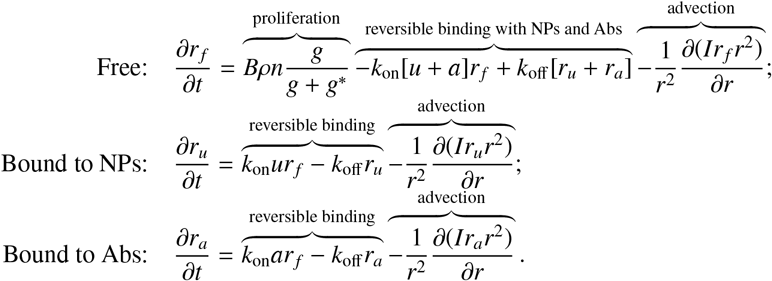

Using the substitution *r* _*f*_ = ρ *f* _*f*_ *n*, as well as analogous substitutions for the other receptor concentrations, and canceling ρ, we obtain:

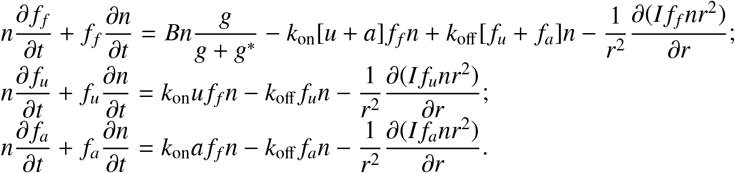

Next, using the equation for the density of proliferating tumor cells (1) and canceling *n*, we obtain:

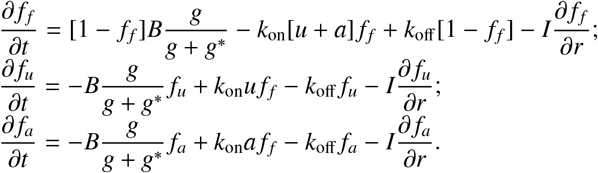

There is no need to consider the equation for *f*_*a*_ explicitly, since it is always equal to 1 − *f*_*f*_ − *f*_*u*_. The other two equations are included in the model.

## Appendix C. Numerical optimization of nanoparticle distribution

As follows from the model equations, the spatial distribution of radiosensitizing nanoparticles that maximizes tumor cell kill during irradiation is governed by the spatial heterogeneity of tumor cell radiosensitivity. In the model, the local effective radiosensitivity is determined by the glucose concentration and can be written as

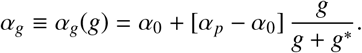

For convenience, we introduce a single variable describing the total local concentration of nanoparticles in tissue, irrespective of whether they are bound to cellular receptors or remain unbound:

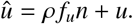

Given a predefined spatial distribution *û*(*r*), the total number of tumor cells killed by an irradiation dose *D* is given by

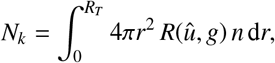

where

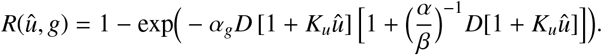

For a fixed irradiation dose *D* and a given glucose distribution *g*(*r*), this formulation makes it possible to determine numerically the optimal spatial distribution *û*(*r*) of a fixed total amount of nanoparticles,

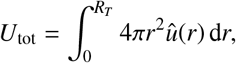

within the tumor volume. For this purpose, we implemented a greedy optimization procedure that repeatedly adds a small concentration increment to *û*(*r*) at those spatial grid points where the derivative ∂*R*(*û*, α_*g*_)/∂*û* attains its largest values. This choice ensures the maximal instantaneous increase in *N*_*k*_ per unit amount of added nanoparticles (note that, due to the model structure, *n* = 1 throughout the entire tumor volume).

Examples of optimal nanoparticle distributions obtained for the model simulation of unconstrained tumor growth, illustrated in Fig. 3a, are shown in Fig. C.1 for different irradiation doses *D* and total nanoparticle amounts *U*_tot_ = 10, 30, and 50 pmol.

**Figure C.1:**
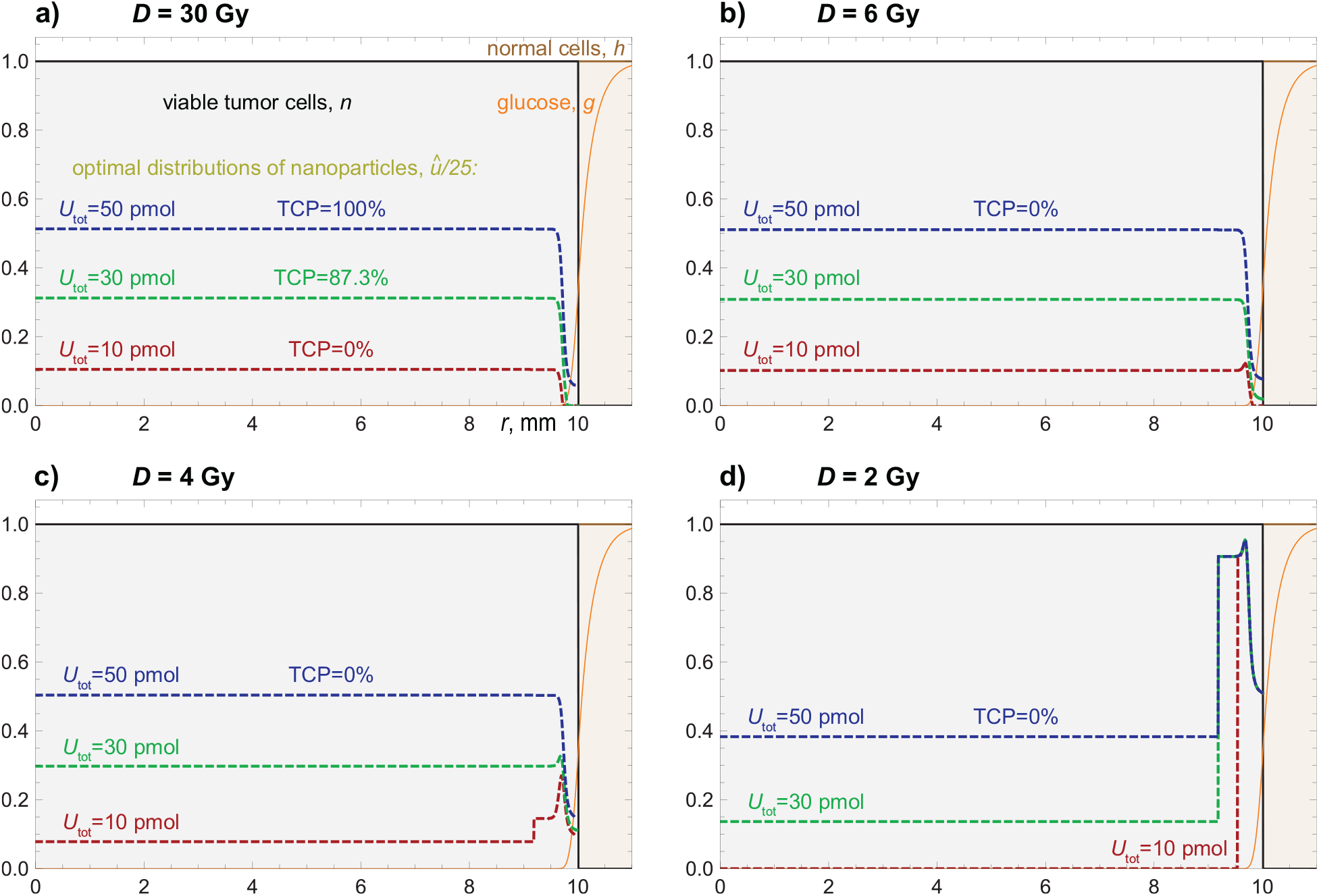
Optimal spatial distributions *û* of fixed total amounts of radiosensitizing nanoparticles (*U*_tot_=10 pmol, 30 pmol, and 50 pmol) that maximize tumor cell kill for different irradiation doses *D*: a) *D* = 30 Gy; b) *D* = 6 Gy; c) *D* = 4 Gy; d) *D* = 2 Gy. Tumor growth is simulated using the basic parameter set from Table 1, and the spatial distributions of viable tumor cells *n*, glucose *g*, and normal cells *h* are identical to those shown in Fig. 3a. TCP denotes the tumor cure probability, defined as TCP = exp(−*N*_*s*_), where *N*_*s*_ is the number of tumor cells surviving irradiation.

As shown in Fig. C.1a, for the highest irradiation dose considered, *D* = 30 Gy (used throughout this study in the model simulations), the optimal nanoparticle distribution preferentially covers the tumor core, which contains the least radiosensitive cells. This implies that the effective irradiation dose *D*_eff_ should be increased in regions of higher cellular radioresistance, in agreement with known analytical results [59]. For *D* = 30 Gy, the use of *U*_tot_ = 50 pmol of nanoparticles is sufficient to ensure tumor cure.

As the irradiation dose is reduced, the structure of the optimal nanoparticle distributions changes. Starting from the lowest values of *U*_tot_, the optimal distributions gradually develop local maxima near the tumor rim. For *U*_tot_ = 10 pmol, such a peak first appears at *D* = 6 Gy (Fig. C.1b) and becomes pronounced at *D* = 4 Gy (Fig. C.1c). At this dose, a similar peak near the tumor boundary is also observed for *U*_tot_ = 30 pmol; however, all depicted optimal profiles still extend into the tumor core, despite yielding zero probability of tumor cure.

For a very low irradiation dose, *D* = 2 Gy, and a small total nanoparticle amount, *U*_tot_ = 10 pmol, the optimal profile *û*(*r*) changes qualitatively: it becomes concentrated exclusively near the tumor rim. In this regime, the optimal distribution resembles that of receptor-bound nanoparticles shortly after intravenous administration rather than that of non-targeted nanoparticles, expected to penetrate deeper into the tumor core.

Taken together, these results indicate that the use of targeted nanoparticles (in the absence of co-administered antibodies capable of redistributing them throughout the tumor volume) can be justified, from the perspective of maximizing tumor cell kill, only in the regime of low irradiation dose and low total nanoparticle amount. Notably, this parameter regime is, by design, insufficient to provide a realistic chance of tumor cure by a single irradiation.

Analytically, it is possible to determine an upper threshold value of the irradiation dose, *D*_th_, at which the structure of the optimal nanoparticle distribution changes qualitatively—from a profile characterized by a high-concentration plateau in the tumor core to one that increasingly emphasizes the tumor rim. The optimal location for an infinitesimal initial increment of nanoparticles can be identified as the spatial position where the derivative

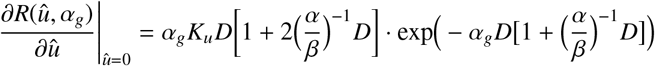

attains its maximum.

Since the glucose level in the tumor core is negligible, the question of whether the first introduced nanoparticles should be placed in the tumor core or near the tumor rim is determined by the sign of the derivative of this expression with respect to α_*g*_(*g*), evaluated at α_*g*_(0) = α_0_:

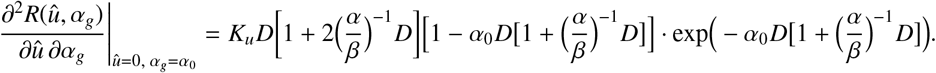

This quantity is positive for sufficiently small values of *D* and negative for its sufficiently large values. In the latter case, it follows that the tumor region with near-zero glucose concentration (the tumor core) represents the optimal location for the first introduced nanoparticles.

The transition between these two regimes occurs when this expression changes sign, which defines the threshold dose *D*_th_ via

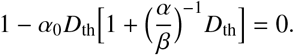

This equation is quadratic in *D*_th_ and has a single positive root,

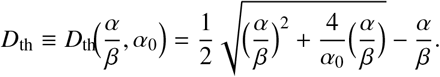

For the basic parameter set used in this study, *D*_th_(10, 0.05) = 10 Gy. This result implies that for irradiation doses close to or exceeding this value, the optimal distribution of even a small amount of nanoparticles, in terms of maximizing tumor cell kill, should preferentially target the relatively radioresistant tumor core.

## Acknowledgements

The graphical abstract was created in BioRender. Kuznetsov, M. (2026). https://BioRender.com/niuba5g

## Data availability

The source code used for numerical simulations, the processed simulation data underlying the figures and population-level analyses, and the Wolfram Mathematica notebook used for data analysis and plotting are available in the associated research dataset [29].

## Declaration of generative AI and AI-assisted technologies in the manuscript preparation process

During the preparation of this work, the authors used ChatGPT (OpenAI) for assistance with language editing, manuscript restructuring, and preparation for submission. The authors reviewed and edited the output as needed and take full responsibility for the content of the published article.

## References

[1] P. Galanakou, T. Leventouri, W. Muhammad, Non-radioactive elements for prompt gamma enhancement in proton therapy, Radiat. Phys. Chem. 196 (2022) 110132.

[2] M. Azarkin, M. Kirakosyan, V. Ryabov, Study of nuclear reactions in therapy of tumors with proton beams, Int. J. Mol. Sci. 24 (17) (2023) 13400.

[3] X. Hu, J. Hu, Y. Pang, M. Wang, W. Zhou, X. Xie, C. Zhu, X. Wang, X. Sun, Application of nano-radiosensitizers in non-small cell lung cancer, Front. Oncol. 14 (2024) 1372780.

[4] C. Bilynsky, N. Millot, A.-L. Papa, Radiation nanosensitizers in cancer therapy—from preclinical discoveries to the outcomes of early clinical trials, Bioeng. Transl. Med. 7 (1) (2022) e10256.

[5] Y.-C. Chuang, P.-H. Wu, Y.-A. Shen, C.-C. Kuo, W.-J. Wang, Y.-C. Chen, H.-L. Lee, J.-F. Chiou, Recent advances in metal-based nanoenhancers for particle therapy, Nanomaterials 13 (6) (2023) 1011.

[6] J.-K. Kim, S.-J. Seo, K.-H. Kim, T.-J. Kim, M.-H. Chung, K.-R. Kim, T.-K. Yang, Therapeutic application of metallic nanoparticles combined with particle-induced x-ray emission effect, Nanotechnology 21 (42) (2010) 425102.

[7] J.-K. Kim, S.-J. Seo, H.-T. Kim, K.-H. Kim, M.-H. Chung, K.-R. Kim, S.-J. Ye, Enhanced proton treatment in mouse tumors through proton irradiated nanoradiator effects on metallic nanoparticles, Phys. Med. Biol. 57 (24) (2012) 8309.

[8] J. C. Polf, L. F. Bronk, W. H. Driessen, W. Arap, R. Pasqualini, M. Gillin, Enhanced relative biological effectiveness of proton radiotherapy in tumor cells with internalized gold nanoparticles, Appl. Phys. Lett. 98 (19) (2011).

[9] S. Zwiehoff, J. Johny, C. Behrends, A. Landmann, F. Mentzel, C. Bäumer, K. Kröninger, C. Rehbock, B. Timmermann, S. Barcikowski, Enhancement of proton therapy efficiency by noble metal nanoparticles is driven by the number and chemical activity of surface atoms, Small 18 (9) (2022) 2106383.

[10] C. Cunningham, M. de Kock, M. Engelbrecht, X. Miles, J. Slabbert, C. Vandevoorde, Radiosensitization effect of gold nanoparticles in proton therapy, Front. Public Health 9 (2021) 699822.

[11] A. L. Popov, D. D. Kolmanovich, N. N. Chukavin, I. V. Zelepukin, G. V. Tikhonowski, A. I. Pastukhov, A. A. Popov, A. E. Shemyakov, S. M. Klimentov, V. A. Ryabov, et al., Boron nanoparticle-enhanced proton therapy: Molecular mechanisms of tumor cell sensitization, Molecules 29 (16) (2024) 3936.

[12] T. Schlathölter, P. Eustache, E. Porcel, D. Salado, L. Stefancikova, O. Tillement, F. Lux, P. Mowat, A. K. Biegun, M.-J. Van Goethem, et al., Improving proton therapy by metal-containing nanoparticles: Nanoscale insights, Int. J. Nanomedicine (2016) 1549–1556.

[13] I. N. Zavestovskaya, A. L. Popov, D. D. Kolmanovich, G. V. Tikhonowski, A. I. Pastukhov, M. S. Savinov, P. V. Shakhov, J. S. Babkova, A. A. Popov, I. V. Zelepukin, et al., Boron nanoparticle-enhanced proton therapy for cancer treatment, Nanomaterials 13 (15) (2023) 2167.

[14] M. Filimonova, D. Kolmanovich, G. Tikhonowski, D. Petrunya, P. Kotelnikova, A. Shitova, O. Soldatova, A. Filimonov, V. Rybachuk, A. Kosachenko, et al., Binary proton therapy of Ehrlich carcinoma using targeted gold nanoparticles, in: Dokl. Biochem. Biophys., Vol. 516, Springer, 2024, pp. 111–114.

[15] M. S. Ryabtseva, M. V. Filimonova, A. S. Filimonov, O. V. Soldatova, A. A. Shitova, V. A. Rybachuk, I. K. Volkova, K. A. Nikolaev, A. O. Kosachenko, S. N. Koryakin, et al., Polyacrylic acid-coated LaB6 nanoparticles as efficient sensitizers for binary proton therapy, Pharmaceutics 17 (4) (2025) 515.

[16] I. N. Zavestovskaya, M. V. Filimonova, A. L. Popov, I. V. Zelepukin, A. E. Shemyakov, G. V. Tikhonowski, M. Savinov, A. S. Filimonov, A. A. Shitova, O. V. Soldatova, et al., Bismuth nanoparticles-enhanced proton therapy: Concept and biological assessment, Mater. Today Nano 27 (2024) 100508.

[17] M. Minnix, L. Li, P. J. Yazaki, A. D. Miller, J. Chea, E. Poku, A. Liu, J. Y. Wong, R. C. Rockne, D. Colcher, et al., TAG-72–targeted α-radionuclide therapy of ovarian cancer using 225Ac-labeled DOTAylated-huCC49 antibody, J. Nucl. Med. 62 (1) (2021) 55–61.

[18] M. Kuznetsov, A. Kolobov, Optimization of size of nanosensitizers for antitumor radiotherapy using mathematical modeling, Int. J. Mol. Sci. 24 (14) (2023) 11806.

[19] G. M. Thurber, M. M. Schmidt, K. D. Wittrup, Factors determining antibody distribution in tumors, Trends Pharmacol. Sci. 29 (2) (2008) 57–61.

[20] B. Menezes, C. Cilliers, T. Wessler, G. M. Thurber, J. J. Linderman, An agent-based systems pharmacology model of the antibody-drug conjugate kadcyla to predict efficacy of different dosing regimens, AAPS J. 22 (2) (2020) 29.

[21] B. Menezes, J. J. Linderman, G. M. Thurber, Simulating the selection of resistant cells with bystander killing and antibody coadministration in heterogeneous human epidermal growth factor receptor 2–positive tumors, Drug Metab. Dispos. 50 (1) (2022) 8–16.

[22] M. Kuznetsov, J. Clairambault, V. Volpert, Improving cancer treatments via dynamical biophysical models, Phys. Life Rev. 39 (2021) 1–48.

[23] T. O. McDonald, Y.-C. Cheng, C. Graser, P. B. Nicol, D. Temko, F. Michor, Computational approaches to modelling and optimizing cancer treatment, Nat. Rev. Bioeng. 1 (10) (2023) 695–711.

[24] K. C. Patra, N. Hay, The pentose phosphate pathway and cancer, Trends Biochem. Sci. 39 (8) (2014) 347–354.

[25] S. K. Stamatelos, E. Kim, A. P. Pathak, A. S. Popel, A bioimage informatics based reconstruction of breast tumor microvasculature with computational blood flow predictions, Microvasc. Res. 91 (2014) 8–21.

[26] I. V. Panyutin, S. A. Holar, R. D. Neumann, I. G. Panyutin, Effect of ionizing radiation on the proliferation of human embryonic stem cells, Sci. Rep. 7 (1) (2017) 43995.

[27] S. J. McMahon, The linear quadratic model: usage, interpretation and challenges, Phys. Med. Biol. 64 (1) (2018) 01TR01.

[28] M. Kuznetsov, V. Adhikarla, E. Caserta, X. Wang, J. E. Shively, F. Pichiorri, R. C. Rockne, Mathematical modeling unveils optimization strategies for targeted radionuclide therapy of blood cancers, Cancer Res. Commun. 4 (11) (2024) 2955–2967.

[29] M. Kuznetsov, A. Kolobov, [dataset] Data and source code for “antibody co-administration robustly improves proton therapy with radiosensitizing nanoparticles: a mathematical modeling study”, Mendeley Data, Version 1 (2026). doi:10.17632/k246r86wgp.1.

[30] W. H. Press, Numerical recipes 3rd edition: The art of scientific computing, Cambridge university press, 2007.

[31] J. Freyer, R. Sutherland, A reduction in the in situ rates of oxygen and glucose consumption of cells in EMT6/Ro spheroids during growth, J. Cell. Physiol. 124 (3) (1985) 516–524.

[32] A. D. Association, et al., Screening for type 2 diabetes, Diabetes Care 27 (suppl 1) (2004) s11–s14.

[33] J. R. Levick, An introduction to cardiovascular physiology, Butterworth-Heinemann, 2013.

[34] A. Bouchoux, H. Roux-de Balmann, F. Lutin, Nanofiltration of glucose and sodium lactate solutions: Variations of retention between single-and mixed-solute solutions, J. Membr. Sci. 258 (1) (2005) 123–132.

[35] U.S. Food and Drug Administration, ELAHERE: Highlights of prescribing information (2022).URL https://www.accessdata.fda.gov/drugsatfda_docs/label/2022/761310s000lbl.pdf

[36] V. Tuchin, A. Bashkatov, E. Genina, Y. P. Sinichkin, N. Lakodina, In vivo investigation of the immersion-liquid-induced human skin clearing dynamics, Tech. Phys. Lett. 27 (6) (2001) 489–490.

[37] J. Casciari, S. Sotirchos, R. Sutherland, Mathematical modelling of microenvironment and growth in EMT6/Ro multicellular tumour spheroids, Cell Prolif. 25 (1) (1992) 1–22.

[38] M. C. Joiner, A. J. van der Kogel, Basic clinical radiobiology, CRC press, 2018.

[39] S. Miotti, P. Facheris, A. Tomassetti, F. Bottero, C. Bottini, F. Ottone, M. I. Colnaghi, M. A. Bunni, D. G. Priest,S. Canevari, Growth of ovarian-carcinoma cell lines at physiological folate concentration: effect on folate-binding protein expression in vitro and in vivo, Int. J. Cancer 63 (3) (1995) 395–401.

[40] O. Ab, K. R. Whiteman, L. M. Bartle, X. Sun, R. Singh, D. Tavares, A. LaBelle, G. Payne, R. J. Lutz, J. Pinkas, et al., IMGN853, a folate receptor-α (FRα)–targeting antibody–drug conjugate, exhibits potent targeted antitumor activity against FRα-expressing tumors, Mol. Cancer Ther. 14 (7) (2015) 1605–1613.

[41] M. B. Kuznetsov, A. V. Kolobov, Transient alleviation of tumor hypoxia during first days of antiangiogenic therapy as a result of therapy-induced alterations in nutrient supply and tumor metabolism–analysis by mathematical modeling, J. Theor. Biol. 451 (2018) 86–100.

[42] L. Zhao, D. Wu, D. Mi, Y. Sun, Radiosensitivity and relative biological effectiveness based on a generalized target model, J. Radiat. Res. 58 (1) (2017) 8–16.

[43] K. Siwowska, S. Haller, F. Bortoli, M. Benesova, V. Groehn, P. Bernhardt, R. Schibli, C. Müller, Preclinical comparison of albumin-binding radiofolates: Impact of linker entities on the in vitro and in vivo properties, Mol. Pharm. 14 (2) (2017) 523–532.

[44] P. M. Smith-Jones, N. Pandit-Taskar, W. Cao, J. O’Donoghue, M. D. Philips, J. Carrasquillo, J. A. Konner, L. J. Old, S. M. Larson, Preclinical radioimmunotargeting of folate receptor alpha using the monoclonal antibody conjugate DOTA–MORAb-003, Nucl. Med. Biol. 35 (3) (2008) 343–351.

[45] D. Alberti, M. van’t Erve, R. Stefania, M. R. Ruggiero, M. Tapparo, S. Geninatti Crich, S. Aime, A quantitative relaxometric version of the ELISA test for the measurement of cell surface biomarkers, Angew. Chem. Int. Ed. 53 (13) (2014) 3488–3491.

[46] I. Zelepukin, A. Yaremenko, E. Petersen, S. Deyev, V. Cherkasov, P. Nikitin, M. Nikitin, Magnetometry based method for investigation of nanoparticle clearance from circulation in a liver perfusion model, Nanotechnology 30 (10) (2019) 105101.

[47] A. A. Popov, I. V. Zelepukin, G. V. Tikhonowski, E. A. Popova-Kuznecova, G. I. Tselikov, A. Al-Kattan, A.-L. Bailly, F. Correard, D. Braguer, M.-A. Esteve, et al., Comparison of pharmacokinetics and biodistribution of laser-synthesized plasmonic Au and TiN nanoparticles, in: J. Phys.: Conf. Ser., Vol. 2058, IOP Publishing, 2021, p. 012004.

[48] J. Kim, C. Bronson, W. L. Hayton, M. D. Radmacher, D. C. Roopenian, J. M. Robinson, C. L. Anderson, Albumin turnover: FcRn-mediated recycling saves as much albumin from degradation as the liver produces, Am. J. Physiol. Gastrointest. Liver Physiol. 290 (2) (2006) G352–G360.

[49] Y. Zhang, W. Du, K. Smuda, R. Georgieva, H. Bäumler, C. Gao, Inflammatory activation of human serum albumin-or ovalbumin-modified chitosan particles to macrophages and their immune response in human whole blood, J. Mater. Chem. B 6 (19) (2018) 3096–3106.

[50] A. K. Iyer, G. Khaled, J. Fang, H. Maeda, Exploiting the enhanced permeability and retention effect for tumor targeting, Drug Discov. Today 11 (17-18) (2006) 812–818.

[51] V. M. Perez-Garcia, L. A. Pérez-Romasanta, Extreme protraction for low-grade gliomas: theoretical proof of concept of a novel therapeutical strategy, Math. Med. Biol. 33 (3) (2016) 253–271.

[52] M. Kuznetsov, A. Kolobov, Spatial optimization of fractionated proton therapy via mathematical modeling, Bull. Lebedev Phys. Inst. 49 (6) (2022) 174–179.

[53] M. Zaider, L. Hanin, Tumor control probability in radiation treatment, Med. Phys. 38 (2) (2011) 574–583.

[54] T. Munro, C. Gilbert, The relation between tumour lethal doses and the radiosensitivity of tumour cells, Br. J. Radiol. 34 (400) (1961) 246–251.

[55] M. D. Mills, R. J. Schulz, Proton-beam therapy: are physicists ignoring clinical realities?, J. Appl. Clin. Med. Phys. 16 (3) (2015) 1.

[56] Z. Chen, M. M. Dominello, M. C. Joiner, J. W. Burmeister, Proton versus photon radiation therapy: A clinical review, Front. Oncol. 13 (2023) 1133909.

[57] E. M. Renkin, Filtration, diffusion, and molecular sieving through porous cellulose membranes, J. Gen. Physiol. 38 (2) (1954) 225–243.

[58] J. Pappenheimer, E. Renkin, L. Borrero, Filtration, diffusion and molecular sieving through peripheral capillary membranes, Am. J. Physiol. 167 (1) (1951) 13–46.

[59] A. Brahme, A. Argren, Optimal dose distribution for eradication of heterogeneous tumors, Acta Oncol. 26 (5) (1987) 377–385.

