## Supplementary material for "Antibody co-administration robustly improves proton therapy with radiosensitizing nanoparticles: a mathematical modeling study": Source code, processed simulation data, and analysis files: Tumor_model_simulation.cpp

```
//*** Source code for the simulations presented in the article
//*** "Antibody co-administration robustly improves proton therapy with radiosensitizing nanoparticles:
//*** a mathematical modeling study"
//*** by Maxim Kuznetsov and Andrey Kolobov
//*** 2026
//
//*** This implementation is intended for Windows and uses several Windows-specific
//*** functions for file and path handling. The code was tested with Dev-C++ 5.11.
//
//*** For improved computational efficiency, the following compiler flags are recommended:
//*** -O3 -march=native -ffast-math -funroll-loops -fomit-frame-pointer -fopenmp-simd
//*** In Dev-C++ 5.11, these can be added via:
//*** Project -> Project Options -> Parameters -> C++ Compiler -> Add
//
//*** Minor modifications may be required when using a different compiler,
//*** development environment, or operating system.

//***********************************************************************************************************************************************************
//********************************* INTRODUCTION ************************************************************************************************************
//***********************************************************************************************************************************************************

//********* Initializing needed components

#include <stdio.h>
#include <windows.h>
#include <ctime>
#include <iostream>
#include <cmath>
#include <conio.h>
#include <fstream>
#include <cstring>

using namespace std;
#define CLAMP01(x) ((x) < 0.0 ? 0.0 : ((x) > 1.0 ? 1.0 : (x)))
#define MIN(a,b) (( (a) < (b) ) ? (a) : (b))
#define MAX(a,b) (( (a) > (b) ) ? (a) : (b))

double output_frequency = 99999999999.0; // how often the distributions of variables will be recorded
double print_frequency = 24.*7.; // how often the current information about simulation will be printed to the screen
bool outputflag = false;
#define rsn_frequency 1. // how often the tumor radius, speed of tumor front and number of tumor cells will be recorded (in hours)

// mode can be set by the folder name, see Begin() 
bool RTmode = true; // irradiation will be done
bool NPmode = true; // nanoparticles will be injected
bool NPlimmode = false; // limiting-case simulation where all receptors are occupied by nanoparticles
bool ATmode = true; // antibodies will be injected

bool RTflag = false;
bool NPflag = false;
bool ATflag = false;

bool InjMadeflag = false;

double RadTreatBegin = 10.; // radius of tumor, at which treatment starts (in mm)
double RadFinal = 13.; // final radius of tumor (in mm)

double Tsurv = 365.*24.; // survival if tumor cell number falls to 0.01
bool SavedInitial = false; // flag of saving pre-treatment distributions

double toutput, tu; 
double survival;

//********* MODEL PARAMETERS

// Fixed parameters

double ag = 0.36; // hydrodynamic radius of glucose
double aa = 5.0; // hydrodynamic radius of antibodies
double au = 10.0; // hydrodynamic radius of nanoparticles

double Dg = 100.0; // glucose diffusion coefficient
double Da = 13.0; // antibodies diffusion coefficient
double Du = 6.5; // nanoparticles diffusion coefficient

double gst = 0.01; // Michaelis constant for glucose consumption rate

double betaalpha = 0.1; // alpha-beta ratio (of radiosensitivity parameters)
double D = 30.0; // radiation dose

double tR = 99999999999.0; // will be defined from the simulation -- 24 hours after reaching RadTreatBegin 

double kon = 1.0; // binding coefficient for receptors with nanoparticles and antibodies
double koff = 0.08; // unbinding coefficient
double Ku = 0.14; // dose enhancement factor of radiosensitizer
double Ca = 0.14; // clearance rate of antibodies
double Au = 10000.; // concentration of nanoparticles in blood upon their injection // 0 or 10000

// Variation in metabolic activity of tumor cells

double B = 0.01; // maximum rate of tumor cell proliferation // 0.01-0.06
double M = B; // dead cell outflow rate
double Q = 1200.0*B; // glucose consumption rate

// Variation in capillary pore size

double mu = 2.0; // capillary pore distribution parameter // 2-4
double Pg; // capillary permeability to glucose // =P(Dg,ag,mu), will be calculated initially // 9.8-37.7
double Pa; // capillary permeability to antibodies // =P(Da,aa,mu), will be calculated initially // 0.26-4.2
double Pu; // capillary permeability to nanoparticles // =P(Du,au,mu), will be calculated initially // 0.006-9.5

// Variation in cell radiosensitivity

double alpha0 = 0.02; // linear parameter of non-proliferating cells radiosensitivity // 0.01-0.1
double alphap = 5.0*alpha0; // linear parameter of proliferating cells radiosensitivity

// Variation in receptor expression levels

double rho = 50.0; // density of specific receptors in the tumor // 50-500

// Variation in clearance rate of nanoparticles

double Cu = 3.0; // clearance rate of nanoparticles // 1-10

// Controlled parameters

double Aa = 1000.; // concentration of antibodies in blood upon their injection // 0, 1000, or 10000
double tNPRT = 999999999.0; // time interval between nanoparticle injection (or just the moment when tumor reaches RadTreatBegin) and irradiation 
							// 24*k, k=0,1,2..., or "infinite" -- for assessment of number of cells that would survive after irradiation

//********* TECHNICAL PARAMETERS

#define size 10000. // radius of a simulation region, in 10^(-2) cm
double R0T = 200.; // initial radius of normal tissue region, in 10^(-2) cm
#define inception_width 2. // radius of initial tumor cell population

#define PI 3.14159265358979323846  // pi
 
//********* SIMULATION PARAMETERS

#define tau_init 0.001
double tau = tau_init; // initial time step
#define dh 0.05 // space step

#define N (int(size/dh)) // number of points in space, minus one
FILE* results;
#define inception_width_int (int)(inception_width/dh) // the size, in cell grids, of initial tumor cell population

//********* COUNTERS

double t = 0.; // main
double ti = 0.; // for output of the distributions of variables
double ts = 0.; // for printing progress on screen
double tr = 0.; // for output of the tumor radius, speed of tumor front and number of tumor cells
int d = 0; // for serial numbers of the files with distributions of variables
int ds = 0; // for serial numbers of printed progress line on screen

//********* MODEL VARIABLES

double n0[N + 1], nr[N + 1], n1[N + 1]; // alive tumor cells
double Nr[N + 1]; // all tumor cells -- for advection equation
double g0[N + 1], G0[N + 1], G1[N + 1], g1[N + 1]; // glucose
double u0[N + 1], ur[N + 1], U0[N + 1], UR[N + 1], U1[N + 1], u1[N + 1]; // free nanoparticles
double a0[N + 1], ar[N + 1], A0[N + 1], AR[N + 1], A1[N + 1], a1[N + 1]; // free antibodies
double ubl, abl; // nanoparticles and antibodies in blood
double ff0[N + 1], ffr[N + 1], ff1[N + 1]; // free receptors
double fu0[N + 1], fur[N + 1], fu1[N + 1]; // receptors bound with nanoparticles
// Receptor "mass" per volume: phi = n * f
double phifr[N + 1];
double phiur[N + 1];

//********* PROCEDURES AND FUNCTIONS

void Begin(); // complex of actions for the beginning, ...
void Cycle_Ending(); // ...end of the main cycle, ...
void End(); // ...and end of the program execution

void Glucose_qs(); // solving the equation for glucose in the quasi-stationary approximation

void Diffusion_NP(); //  solving the equation for diffusion of nanoparticles
void Diffusion_AT(); //  solving the equation for diffusion of antibodies

void TMA(double apr, double* bpr, double cpr, double* F, double* prog,
	double muleft, double hileft, double muright, double hiright); // tridiagonal matrix algorithm
void TMAc(double apr, double bpr, double cpr, double* F, double* prog,
	double muleft, double hileft, double muright, double hiright);
	
void EulerAdv(); // Euler+advection method
double MMg[N + 1]; // auxiliary variable for proliferation term

void PrepareRT(); // Prepare the cell kill due to RT
void AssessRT(); // Estimate the remaining number of cells
void PerformRT();
double RTcoef[N + 1], Deff[N + 1], alpha_eff[N + 1]; // auxiliary arrays
double nRTsurv[N + 1];

void Profile_output(); // regular output of distributions of variables
void PrintProgress(); // print progress on screen
void WriteProfile(FILE* out);  // output of distributions of variables before treatment
void SavePreTreatmentProfile(double t_current); 

void Radius_estimation();
void Speed_estimation();
void Radius_Speed_CellNumber_output();

//********* AUXILIARY VARIABLES

FILE* inputInitial;
double lenmmprev, lenmm, nIn, gIn, uIn, ffIn, fuIn, aIn;

////// for glucose quasistationary solution
double alpha_g[N + 1];
double muleftg = 0.0;
double hileftg = 0.5 / 1.5;
double murightg = 0.0;
double hirightg;
double norm_g;
double norm_min = 1.e-6; // allowed error during iterative solution
double aprg = 1.;
double cprg = 1.;
double bprg[N + 1];
double Fg[N + 1];
double progg[N + 1];
double pp[N + 1], qq[N + 1];
double kg = Dg / (dh*dh);
int kqs; // counter of number of iterations
int k_crit = 10000; // maximum number of iterations
double gstat; // stationary values in absence of tumor

////// for nanoparticles diffusion
double ku = 2 * dh * dh / (tau * Du);
double apru = 1.;
double bpru = 2. + ku;
double cpru = 1.;
double muleftu = 0.;
double hileftu = 0.5 / 1.5;
double murightu = 0.;
double hirightu;	
double Fu[N + 1];
double progu[N + 1];

////// for antibodies diffusion
double ka = 2 * dh * dh / (tau * Da);
double apra = 1.;
double bpra = 2. + ka;
double cpra = 1.;
double mulefta = 0.;
double hilefta = 0.5 / 1.5;
double murighta = 0.;
double hirighta;
double Fa[N + 1];
double proga[N + 1];

int NN = inception_width_int; // number of points in space where tumor cells are
int NH = (int(R0T / dh)); // number of point in space where tumor and normal cells are
double hNN = 0.; // the variable size of the last grid element, NN, where tumor cells are
double hNH = 0.; // the variable size of the last grid element, NH, where normal cells are

////// for advection
double dN;
double V[N + 1], Vvar[N + 1], dV[N + 1];
// radii of cell centers: Rmid[i] = (i + 0.5)*dh, and their reciprocals
double Rmid[N + 1];
double Rmid_inv[N + 1];
int i0[N + 1];
int Ni0;
int sign;
int NNold;
double hNNold; // for solution of cells equations
double RNnew;
double RHnew;

////// for tumor radius, speed of tumor front and number of tumor cells
double RadTis; // current whole tissue radius, in mm
double Rad; // current tumor radius, in mm
double Rad_prev; // previously measured tumor radius, in mm
double T_prev; // time of previous measurement of tumor radius
double Vfr; // tumor front speed
double NTC; // total number of tumor cells
double SURV; // number of surviving cells if RT takes place now
double SURVmin;
double tSURVmin;

////// for work with files for distribution of variables
char File_name1[100], num[10], radiusTum[8], radiusTis[8];

inline double fast_cbrt(double x)
{
    return cbrt(x);
}

// Permeability calculation

// ===========================
// Lognormal PDF
// ===========================
double L_pdf(double r, double mu, double sigma = 0.5)
{
    if (r <= 0.0) return 0.0;
    double x = (std::log(r) - mu) / sigma;
    return (1.0 / (r * sigma * std::sqrt(2.0 * M_PI))) * std::exp(-0.5 * x * x);
}

// ===========================
// Renkin equation
// ===========================
double Dprime(double D, double a, double r)
{
    if (r <= a) return 0.0;
    double x = a / r;
    return D * (1.0 - 2.1 * x + 2.09 * x * x * x - 0.95 * std::pow(x, 5));
}

// ===========================
// Integrand for permeability
// ===========================
double Integrand(double D, double a, double r, double mu)
{
    if (r <= a) return 0.0;

    double dpr = Dprime(D, a, r);
    double area = (r - a) * (r - a);

    double L = L_pdf(r, mu);

    return L * area * dpr;
}

// ===========================
// P(D, a, mu) permeability
// ===========================
// Uses numerical integration from r=0 to r=1000 nm (more than enough)
double Permeability(double D, double a, double mu)
{
    const double norm = 1.0 / (50.0 * std::pow(mu, 4)); // normalization
    const double rmax = 1000.0;  // nm � enough because pdf decays rapidly
    const int K = 20000;         // resolution
    const double dr = rmax / K;

    double integral = 0.0;

    // Trapezoidal integration
    for (int i = 1; i < K; i++)
    {
        double r = i * dr;
        integral += Integrand(D, a, r, mu);
    }

    // Add edges
    integral += 0.5 * Integrand(D, a, 0.0, mu);
    integral += 0.5 * Integrand(D, a, rmax, mu);

    integral *= dr;

    return norm * integral;
}

//***********************************************************************************************************************************************************
//********************************* MAIN CYCLE **************************************************************************************************************
//***********************************************************************************************************************************************************

int main()
{
	Begin();

	while (Rad <= RadFinal && NTC > 0.01)
	{		
		t = t + tau;
		Radius_estimation();
		Speed_estimation();
		
		Glucose_qs();
		EulerAdv();		
		
		if (NPflag)
			Diffusion_NP();
			
		if (ATflag)
			Diffusion_AT();
			
		if (t >= tr - 0.00001) // small epsilon to avoid floating-point roundoff issues
		{
			if(Rad > RadTreatBegin)
			{
				PrepareRT();
				AssessRT();
			}
			Radius_Speed_CellNumber_output();
		}
		
		if (outputflag && t >= ti - 0.00001) 
			Profile_output();
			
		if (t >= ts - 0.00001) 
			PrintProgress();

		Cycle_Ending();
		
		if (Rad > RadTreatBegin - dh && !SavedInitial)
		{
		    SavePreTreatmentProfile(t); 
		    SavedInitial = TRUE;	
		}

		if (Rad > RadTreatBegin && tR > t + tNPRT + 0.1 && !InjMadeflag)
		{
			if (outputflag)
				toutput = t;
			
			tu = t;
			tR = t + tNPRT; 
			survival = t;
			SURVmin = 2.*NTC;
			
			if (NPmode)
			{
				ubl = ubl + Au;
				NPflag = true;
				printf("\nTime of nanoparticles injection: t = %.1f days\n", t/24.);	
			}
				
			if (ATmode)
			{	
				abl = abl + Aa;
				ATflag = true;
				printf("\nAntibodies are co-injected\n");
			}
			
			if (RTmode)
			{	
				if (outputflag)
				{
					output_frequency = 1.;
					ti = t + output_frequency;
				}
				if (tNPRT < 99999999.0)
				printf("\nIrradiation will be performed in %.1f hours, now t = %.1f days\n", tNPRT, t/24.);	
			}
			
			InjMadeflag = true;
		}
		
		if (RTmode && t > tR && !RTflag)
		{
			RTflag = true;
			PrepareRT();
			PerformRT();
			
			printf("\nIrradiation is performed\n");	
		}
		
		if (NPflag)
		    ubl *= (1.0 - Cu * tau);
		
		if (ATflag)
		    abl *= (1.0 - Ca * tau);
		    
	/*	if (outputflag && t > toutput + 280.)
		{
			output_frequency = 7.*24.;
			ti = t + output_frequency;
			toutput = 99999999999.;			
		}*/

	}
	
	End();
}

//***********************************************************************************************************************************************************
//********************************* BEGINNING OF SIMULATION *************************************************************************************************
//***********************************************************************************************************************************************************

void Begin()
{
	// parameters can be set in the name of the folder where .exe file is located
	// e.g., "mode=AT output B=0.01 mu=2 alpha0=0.02 rho=50 Cu=3 tNPRT=24 Aa=1 kon=0"
	// modes can be Free, RT, NP, limNP, AT

	char buffer[MAX_PATH];
	GetModuleFileNameA(NULL, buffer, MAX_PATH);

	string str(buffer);

	// mode Free
	size_t pos_free = str.find("mode=Free");
	if (pos_free != -1)
	{
		RTmode = false; // irradiation will not be done
		NPmode = false; // nanoparticles will not be injected
		NPlimmode = false; // no limit simulation where all receptors have nanoparticles attached
		ATmode = false; // antibodies will not be injected
		
		cout << "Simulating free growth\n";
	}

	// mode RT
	size_t pos_RT = str.find("mode=RT");
	if (pos_RT != -1)
	{
		RTmode = true; // irradiation will be done
		NPmode = false; // nanoparticles will not be injected
		NPlimmode = false; // no limit simulation where all receptors have nanoparticles attached
		ATmode = false; // antibodies will not be injected
		
		cout << "Simulating RT, no nanoparticles\n";
	}

	// mode NP
	size_t pos_NP = str.find("mode=NP");
	if (pos_NP != -1)
	{
		RTmode = true; // irradiation will be done
		NPmode = true; // nanoparticles will be injected
		NPlimmode = false; // no limit simulation where all receptors have nanoparticles attached
		ATmode = false; // antibodies will not be injected
		
		cout << "Simulating RT with nanoparticles, no antibodies\n";
	}	
	
	// mode limNP
	size_t pos_limNP = str.find("mode=limNP");
	if (pos_limNP != -1)
	{
		RTmode = true; // irradiation will be done
		NPmode = false; // nanoparticles will not be injected
		NPlimmode = true; // limit simulation where all receptors have nanoparticles attached
		ATmode = false; // antibodies will not be injected
		
		cout << "Simulating RT in limiting case of all receptors occupied by NPs\n";
	}	

	// mode AT
	size_t pos_AT = str.find("mode=AT");
	if (pos_AT != -1)
	{
		RTmode = true; // irradiation will be done
		NPmode = true; // nanoparticles will be injected
		NPlimmode = false; // no limit simulation where all receptors have nanoparticles attached
		ATmode = true; // antibodies will be injected
		
		cout << "Simulating RT with nanoparticles and antibodies\n";
	}	
	
	// output
	size_t pos_output = str.find("output");
	if (pos_output != -1)
	{
		outputflag = true;
		output_frequency = 7.*24.;
		
		cout << "\nProfiles will be recorded\n";
	}
	
	// B
	size_t pos_B = str.find("B=");
	if (pos_B != -1)
	{
		string str_B = str.substr(pos_B + 2);
		B = atof(str_B.c_str());
	}

	cout << "B=" << B << "\n";	
	
	// mu
	size_t pos_mu = str.find("mu=");
	if (pos_mu != -1)
	{
		string str_mu = str.substr(pos_mu + 3);
		mu = atof(str_mu.c_str());
	}

	cout << "mu=" << mu << "\n";

	if (RTmode == true)
	{
		// alpha0
		size_t pos_alpha0 = str.find("alpha0=");
		if (pos_alpha0 != -1)
		{
			string str_alpha0 = str.substr(pos_alpha0 + 7);
			alpha0 = atof(str_alpha0.c_str());
		}

		cout << "alpha0=" << alpha0 << "\n";
	
		// rho
		size_t pos_rho = str.find("rho=");
		if (pos_rho != -1)
		{
			string str_rho = str.substr(pos_rho + 4);
			rho = atof(str_rho.c_str());
			
			if (rho == 0.)
				kon = 0.; // since rho does not explicitly appear in the equations for receptor fractions
		}
		
		cout << "rho=" << rho << "\n";			
	
		// tNPRT
		size_t pos_tNPRT = str.find("tNPRT=");
		if (pos_tNPRT != -1)
		{
			string str_tNPRT = str.substr(pos_tNPRT + 6);
			tNPRT = atof(str_tNPRT.c_str());
		}
	
		cout << "tNPRT=" << tNPRT << "\n";	
	}
	
	if (NPmode == true)
	{
		// Cu
		size_t pos_Cu = str.find("Cu=");
		if (pos_Cu != -1)
		{
			string str_Cu = str.substr(pos_Cu + 3);
			Cu = atof(str_Cu.c_str());
		}
	
		cout << "Cu=" << Cu << "\n";
		
		// kon
		size_t pos_kon = str.find("kon=");
		if (pos_kon != -1)
		{
			string str_kon = str.substr(pos_kon + 4);
			kon = atof(str_kon.c_str());
		}
	
		cout << "kon=" << kon << "\n";
	}
	
	if (ATmode == true)
	{
		// Aa
		size_t pos_Aa = str.find("Aa=");
		if (pos_Aa != -1)
		{
			string str_Aa = str.substr(pos_Aa + 3);
			Aa = atof(str_Aa.c_str());
		}
	
		cout << "Aa=" << Aa << "\n";	
	}
	
	cout << "\n";
	
	// calculating permeability values
	Pg = Permeability(Dg, ag, mu);
	Pa = Permeability(Da, aa, mu);
	Pu = Permeability(Du, au, mu);

	// "volumes" of grid elements
	for (int i = 0; i <= N; i++)
		V[i] = (i + 1)*dh*(i + 1)*dh*(i + 1)*dh - i*dh*i*dh*i*dh;

    // pre-calculate the radii of cell centers and their reciprocals
    for (int i = 0; i <= N; i++)
    {
        Rmid[i]     = ((double)i + 0.5) * dh;
        Rmid_inv[i] = 1.0 / Rmid[i];
    }
		
	//********* CREATING FILE FOR TUMOR RADIUS, FRONT SPEED AND CELL NUMBER
	if ((results = fopen("results.txt", "wt")) == NULL)
		fprintf(stderr, "Cannot open file results.txt\n\n");

	fprintf(results, "t(h)	RTum	Speed 	TumCells 	SURV 		RTis(mm) 	ubl 		abl\n\n");
	fflush(results);

	//********* INITIAL CONDITIONS
	lenmmprev = 0.;

	// trying to find any Initial*.txt
	char initialName[MAX_PATH];
	HANDLE hFind;
	WIN32_FIND_DATAA ffd;
	
	hFind = FindFirstFileA("Initial*.txt", &ffd);
	if (hFind != INVALID_HANDLE_VALUE)
	{
	    // found one
	    strcpy(initialName, ffd.cFileName);
	    FindClose(hFind);
	
	    // we try to extract the time from the name if it contains "t="
	    double t_init = 0.0;
	    char* p = strstr(initialName, "t=");
	    if (p != NULL)
	    {
	        t_init = atof(p + 2); // after 't='
	        t = t_init;           // setting initial time in the model
	        ti = t + output_frequency;
	        tr = t + rsn_frequency;
	        ts = t;
			d = 0;
			ds = 0;
	    }
	
	    if ((inputInitial = fopen(initialName, "rt")) != NULL)
	    {
	        cout << "Reading initial conditions from " << initialName << "\n";
	        NN = 0;
	        NH = 0;
	
	        char* word;
	        int iScan;
	        word = new char[512];
	        fgets(word, 512, inputInitial);
	        fgets(word, 512, inputInitial);
	
	        do {
			    if (!fgets(word, 512, inputInitial))
			        break; 
			
			    iScan = sscanf(word, "%lf%lf%*lf%*lf%lf%lf%lf%lf%lf", &lenmm, &nIn, &gIn, &uIn, &ffIn, &fuIn, &aIn);
			    if (iScan < 3)
			        break; 
			
			    lenmmprev = lenmm;
			
			    n0[NN] = nIn;
			    g0[NN] = gIn;
			    u0[NN] = uIn;
			    ff0[NN] = ffIn;
			    fu0[NN] = fuIn;
			    a0[NN] = aIn;
			    NN++;
			} while (nIn > 0.);

	        NN--;
	        NN--;
	        hNN = 10. * lenmmprev - (1. * NN) * dh;
	        NH = NN + 1;
	        do {
			    if (!fgets(word, 512, inputInitial))
			        break;
			
			    iScan = sscanf(word, "%lf%lf%*lf%*lf%lf%lf%lf%lf%lf", &lenmm, &nIn, &gIn, &uIn, &ffIn, &fuIn, &aIn);
			    if (iScan < 3)
			        break;
			
			    lenmmprev = lenmm;
			    n0[NH] = nIn;
			    g0[NH] = gIn;
			    u0[NH] = uIn;
			    ff0[NH] = ffIn;
			    fu0[NH] = fuIn;
			    a0[NH] = aIn;
			    NH++;
			} while (!feof(inputInitial));

	        NH--;
	        NH--;
	        hNH = 10. * lenmmprev - (1. * NH) * dh;
	        fclose(inputInitial);
	        SavedInitial = TRUE;
	    }
	    else
	    {
	        fprintf(stderr, "Cannot open %s, fallback to default IC\n", initialName);
	        goto default_IC;
	    }
	}
	else
	{
		default_IC:
	    //********* INITIAL CONDITIONS
	    for (int i = 0; i <= NH; i++)
	    {
	        n0[i] = 0.;
	        g0[i] = 1.;
		    u0[i] = 0.;
		    ff0[i] = 0.;
		    fu0[i] = 0.;
		    a0[i] = 0.;
	    }
	
	    for (int i = 0; i <= NN; i++)
	    {
	        n0[i] = 1.;
	        ff0[i] = 1.;
	    }
	}

	for (int i = 0; i <= N; i++) // this is made for easy output of the initial conditions
	{
		n1[i] = n0[i];
		g1[i] = g0[i];
		u1[i] = u0[i];
		ff1[i] = ff0[i];
		fu1[i] = fu0[i];
		a1[i] = a0[i];
	}

	Glucose_qs(); // replaced with quasistationary values
	gstat = g0[NH]; // stationary values in absence of tumor

	Radius_estimation();
	
	T_prev  = t;
	Rad_prev = Rad;
	Vfr = 0.;

	//********* COUNTING TUMOR CELLS		
	NTC = 0.;

	for (int i = 0; i <= NN - 1; i++)
		NTC = NTC + V[i] * n1[i];

	NTC = NTC + ((NN*dh + hNN)*(NN*dh + hNN)*(NN*dh + hNN) - NN*dh*NN*dh*NN*dh) * n1[NN];

	NTC = 4. * PI * 300. * NTC / 3.;

	Radius_Speed_CellNumber_output();
	
	if (outputflag)
		Profile_output();
}

//***********************************************************************************************************************************************************
//********************************* OUTPUT OF DISTRIBUTIONS OF VARIABLES ************************************************************************************
//***********************************************************************************************************************************************************

void WriteProfile(FILE* out)
{
    fprintf(out, "Len(mm)\tn\t\tn+m\tn+m+h\tg\t\tu\t\tff\tfu\ta\t\t\n\n");

    // 1) Inside tumor: 0 .. NN-1
    for (int i = 0; i <= NN - 1; i++)
    {
        double len_mm = ((double)i + 0.5) * dh / 10.0;
        double nm = 1.0;    // n+m (tumor occupancy)
        double h  = 0.0;    // normal cells

        fprintf(out, "%.4f\t%.5e\t%g\t%g\t%.5e\t%.5e\t%.5f\t%.5f\t%.5e\n",
                len_mm,
                n1[i],
                nm,
                nm + h,    // here = 1.0
                g1[i],
				u1[i],
				ff1[i],
				fu1[i],
				a1[i]);
    }

    // 2) Boundary point NN (irregular width hNN)
    {
        double len_mm = ((double)NN * dh + hNN) / 10.0;

        double g_interp = g1[NN - 1] +
                          (g1[NN + 1] - g1[NN - 1]) *
                          ((dh / 2.0) + hNN) / (2.0 * dh);

        double u_interp = u1[NN - 1] +
                          (u1[NN + 1] - u1[NN - 1]) *
                          ((dh / 2.0) + hNN) / (2.0 * dh);

        double a_interp = a1[NN - 1] +
                          (a1[NN + 1] - a1[NN - 1]) *
                          ((dh / 2.0) + hNN) / (2.0 * dh);

        double nm = 1.0;  // still tumor region (last shell)
        double h  = 0.0;

        fprintf(out, "%.4f\t%.5e\t%g\t%g\t%.5e\t%.5e\t%.5f\t%.5f\t%.5e\n",
                len_mm,
                n1[NN],
                nm,
                nm + h,
                g_interp,
				u_interp,
				ff1[NN],
				fu1[NN],
				a_interp);
    }

    // 3) Outside tumor: NN+1 .. NH-1
    for (int i = NN + 1; i <= NH - 1; i++)
    {
        double len_mm = ((double)i + 0.5) * dh / 10.0;
        double nm = 0.0;
        double h  = 1.0;

        fprintf(out, "%.4f\t%.5e\t%g\t%g\t%.5e\t%.5e\t%.5f\t%.5f\t%.5e\n",
                len_mm,
                n1[i],       // ~0
                nm,
                nm + h,      // = 1.0
                g1[i],
				u1[i],
				0.0,
				0.0,
				a1[i]);
    }

    // 4) Final boundary NH (irregular width hNH)
    {
        double len_mm = ((double)NH * dh + hNH) / 10.0;
        double nm = 0.0;
        double h  = 1.0;

        fprintf(out, "%.4f\t%.5e\t%g\t%g\t%.5e\t%.5e\t%.5f\t%.5f\t%.5e\n",
                len_mm,
                n1[NH],
                nm,
                nm + h,
                g1[NH],
				u1[NH],
				0.0,
				0.0,
				a1[NH]);
    }
}

void SavePreTreatmentProfile(double t_current)
{
    char fname[256];
    sprintf(fname, "PreTreatment t=%.6f.txt", t_current);

    FILE* out = fopen(fname, "wt");
    if (!out)
    {
        fprintf(stderr, "Cannot open pretreatment file %s\n", fname);
        return;
    }
    setvbuf(out, NULL, _IOFBF, 1 << 20);

    WriteProfile(out);

    fclose(out);

    printf("\nSaved pretreatment profile: %s\n", fname);
}

void Profile_output()
{
    //********* CREATING A FILENAME WITH SERIAL NUMBER

    _itoa(d, num, 10);

    if (d < 10)
        strcpy(File_name1, "res-d0000");
    else if (d < 100)
        strcpy(File_name1, "res-d000");
    else if (d < 1000)
        strcpy(File_name1, "res-d00");
    else if (d < 10000)
        strcpy(File_name1, "res-d0");
    else
        strcpy(File_name1, "res-d");

    strcat(File_name1, num);

    // DAY 
    double day = t / 24.0;
    char dayStr[16];
    sprintf(dayStr, "%.1f", day);

    strcat(File_name1, "_day=");
    strcat(File_name1, dayStr);

    // TUMOR RADIUS 
    if (Rad < 1.)
    {
        _gcvt(Rad, 1, radiusTum);
        if (radiusTum[1] == '\0')
            radiusTum[1] = '0';
    }
    else if (Rad < 10.)
    {
        _gcvt(Rad, 2, radiusTum);
        if (radiusTum[2] == '\0')
            radiusTum[2] = '0';
    }
    else
    {
        _gcvt(Rad, 3, radiusTum); // in mm
        if (radiusTum[3] == '\0')
            radiusTum[3] = '0';
    }

    strcat(File_name1, "_RadTum=");
    strcat(File_name1, radiusTum);

    strcat(File_name1, ".txt");

    // OUTPUT
    FILE* out;
    if ((out = fopen(File_name1, "wt")) == NULL)
    {
        fprintf(stderr, "Cannot open file\n");
        return;
    }
    setvbuf(out, NULL, _IOFBF, 1 << 20);

    WriteProfile(out);

    fclose(out);

    // switch counters
    ti += output_frequency;
    d++;
}

void PrintProgress()
{
    // PRINTING ON SCREEN
    printf("#%i, t = %.1f days, radius = %.3f mm, speed = %.3f mm/week, cell number = %.3e\n",
           ds, t/24., Rad, Vfr, NTC);
           
    // switch counters
    ts += print_frequency;
    ds++;
}


//***********************************************************************************************************************************************************
//********************************* ESTIMATION OF RADIUS AND SPEED ******************************************************************************************
//***********************************************************************************************************************************************************

//********* ESTIMATION OF RADIUS

void Radius_estimation()
{
	Rad = (NN * dh + hNN) / 10.;
	RadTis = (NH * dh + hNH) / 10.;
}

//********* ESTIMATION OF TUMOR FRONT SPEED

void Speed_estimation()
{
	if (t - T_prev > 5. || (abs(Rad - Rad_prev) > 0.2 && 10. * Rad > R0T))
	{
		Vfr = 7. * 24. * (Rad - Rad_prev) / (t - T_prev); // in mm/week
		T_prev = t;
		Rad_prev = Rad;
	}
}

//***********************************************************************************************************************************************************
//********************************* OUTPUT OF THE TUMOR RADIUS, SPEED OF TUMOR FRONT AND NUMBER OF TUMOR CELLS **********************************************
//***********************************************************************************************************************************************************

void Radius_Speed_CellNumber_output()
{
	tr = tr + rsn_frequency;

	//********* COUNTING TUMOR CELLS

	NTC = 0.;

	for (int i = 0; i <= NN - 1; i++)
		NTC = NTC + V[i] * n1[i];

	NTC = NTC + ((NN*dh + hNN)*(NN*dh + hNN)*(NN*dh + hNN) - NN*dh*NN*dh*NN*dh) * n1[NN];

	NTC = 4. * PI * 300. * NTC / 3.;
	
	//********* OUTPUT	
	fprintf(results, "%.1f\t%.5f\t%.5f\t%.3e\t%.3e\t%.5f\t%.3e\t%.3e\t\n", t, Rad, Vfr, NTC, SURV, RadTis, ubl, abl);
	fflush(results);
}

//***********************************************************************************************************************************************************
//********************************* SOLUTION OF KINETIC TERMS ***********************************************************************************************
//***********************************************************************************************************************************************************

//********* EULER PROCEDURE + ADVECTION

void EulerAdv()
{
    // Try to slowly restore time step up to tau_init
    tau = MIN(2.0 * tau, tau_init);
    ku = 2 * dh * dh / (tau * Du);
    ka = 2 * dh * dh / (tau * Da);

tryAgain:

    // --- 1) Reaction terms ---
    for (int i = 0; i <= NN; i++)
    {
        MMg[i] = g0[i] / (g0[i] + gst);

        // Ensure values stay in [0,1]
        n0[i] = CLAMP01(n0[i]);

        // nr = new density of alive tumor cells
        nr[i] = n0[i] + tau * (B * n0[i] * MMg[i]);

        // Nr = alive + dead = updated total tumor density
        Nr[i] = nr[i] + (1.0 - n0[i]) - tau * M * (1.0 - n0[i]);
    }
    
    if(NPflag)
    {
    	for (int i = 0; i <= NN; i++)
        {
	        ffr[i] = ff0[i] + tau * ((1.0 - ff0[i]) * B * MMg[i] - kon * (u0[i] + a0[i]) * ff0[i] + koff*(1.0 - ff0[i]));
	        fur[i] = fu0[i] + tau * (- B * MMg[i] * fu0[i] + kon * u0[i] * ff0[i] - koff * fu0[i]);   
	    }    	
    	
    	for (int i = 0; i <= NN - 1; i++) 
	 	  	ur[i] = u0[i] + tau * (- kon * rho * ff0[i] * n0[i] * u0[i] + koff * rho * fu0[i] * n0[i]);
	 	  	
	 	 // Geometry at the tumor boundary (cell NN and NN+1)
        const double R_NN    = NN * dh;
        const double R_NNp1  = (NN + 1) * dh;
        const double R_NNh   = R_NN + hNN;

        const double R_NN3   = R_NN   * R_NN   * R_NN;
        const double R_NNp13 = R_NNp1 * R_NNp1 * R_NNp1;
        const double R_NNh3  = R_NNh  * R_NNh  * R_NNh;

        const double full_shell = R_NNp13 - R_NN3;   // Full volume of shell [NN, NN+1]
        const double tum_shell  = R_NNh3  - R_NN3;   // Portion filled by tumor
        const double out_shell  = R_NNp13 - R_NNh3;  // Outer portion

        const double ratio_tum = tum_shell / full_shell;
        const double ratio_out = out_shell / full_shell;
        
        ur[NN] = u0[NN] + tau * (Pu * (ubl - u0[NN]) * ratio_out - kon * rho * ff0[NN] * n0[NN] * u0[NN] * ratio_tum + koff * rho * fu0[NN] * n0[NN] * ratio_tum);
        
    	for (int i = NN + 1; i <= NH; i++) 
	 	  	ur[i] = u0[i] + tau * (Pu * (ubl - u0[i]));
	}
	
    if(ATflag)
    {    	
    	for (int i = 0; i <= NN - 1; i++) 
	 	  	ar[i] = a0[i] + tau * (- kon * rho * ff0[i] * n0[i] * a0[i] + koff * rho * (1.0 - ff0[i] - fu0[i]) * n0[i]);
	 	  	
	 	 // Geometry at the tumor boundary (cell NN and NN+1)
        const double R_NN    = NN * dh;
        const double R_NNp1  = (NN + 1) * dh;
        const double R_NNh   = R_NN + hNN;

        const double R_NN3   = R_NN   * R_NN   * R_NN;
        const double R_NNp13 = R_NNp1 * R_NNp1 * R_NNp1;
        const double R_NNh3  = R_NNh  * R_NNh  * R_NNh;

        const double full_shell = R_NNp13 - R_NN3;   // Full volume of shell [NN, NN+1]
        const double tum_shell  = R_NNh3  - R_NN3;   // Portion filled by tumor
        const double out_shell  = R_NNp13 - R_NNh3;  // Outer portion

        const double ratio_tum = tum_shell / full_shell;
        const double ratio_out = out_shell / full_shell;
        
        ar[NN] = a0[NN] + tau * (Pa * (abl - a0[NN]) * ratio_out - kon * rho * ff0[NN] * n0[NN] * a0[NN] * ratio_tum + koff * rho * (1.0 - ff0[NN] - fu0[NN]) * n0[NN] * ratio_tum);
        
    	for (int i = NN + 1; i <= NH; i++) 
	 	  	ar[i] = a0[i] + tau * (Pa * (abl - a0[i]));
	}

    // --- 2) Advection / redistribution of volume ---
    // Special handling of NN==0: tumor fits into one computational cell

    if (NN == 0)
    {
        // === SPECIAL CASE: tumor occupies a single grid cell ===

        // Old radius Rold = hNN because NN = 0
        double Rold  = hNN;
        double Rold3 = Rold * Rold * Rold;

        // Volume of the spherical shell corresponding to this cell
        double Vold = Rold3;   // inner radius is zero here

        // dN = volume change determined by deviation of Nr from 1
        dN = (Nr[0] - 1.0) * Vold;

        // Save old width for stability check
        hNNold = hNN;

        // New radius of the tumor from volume conservation
       	double arg = Rold3 + dN;
		RNnew = (arg <= 0.0) ? 0.0 : fast_cbrt(arg);

        hNN = RNnew;  // for NN=0 this is the full radius

        // CFL-like stability check
        double dh_diff = hNN - hNNold;
        if (dh_diff < 0.0) dh_diff = -dh_diff;
        if (dh_diff > dh)
        {
            tau *= 0.5;
            goto tryAgain;
        }

        // No fluxes inside a single-cell domain, only reaction applies
        n1[0] = nr[0] / Nr[0];
		n1[0] = CLAMP01(n1[0]);
		
		if (NPflag || ATflag)
		{
		    ff1[0] = ffr[0];
		    fu1[0] = fur[0];
		}
		else
		{
		    ff1[0] = 1.0;
		    fu1[0] = 0.0;
		}

        // Enforce bounds
        n1[0] = CLAMP01(n1[0]);
        
		// If new boundary exceeds one grid cell, add a new one
        while (hNN > dh)
        {
            NN  = NN + 1;
            n1[NN] = n1[NN - 1];  // copy boundary value
            ff1[NN] = ff1[NN - 1]; 
            fu1[NN] = fu1[NN - 1]; 
            hNN   = hNN - dh;
        }
        
		for (int i = 0; i <= NN; i++)
       		n1[i] = CLAMP01(n1[i]);
    }
    else
    {
        // === GENERAL CASE: NN >= 1 � full redistribution scheme ===

        // Precompute geometry for current NN, hNN
        double R_NN    = NN * dh;
        double R_NN_h  = R_NN + hNN;
        double R_NN3   = R_NN * R_NN * R_NN;
        double R_NN_h3 = R_NN_h * R_NN_h * R_NN_h;

        dN    = 0.0;
        Ni0   = 1;
        i0[0] = -1;

        for (int i = 0; i <= NN; i++)
            dV[i] = 0.0;

        // First cell (i=0)
        dV[0] = dN;
        dN   += (Nr[0] - 1.0) * V[0];

        // Interior cells (1 .. NN-1)
        for (int i = 1; i <= NN - 1; i++)
        {
            dV[i] = dN;

            // Stability check: cannot shift more than neighbor volume
            double dv_abs = dV[i];
            if (dv_abs < 0.0) dv_abs = -dv_abs;
            if (dv_abs > V[i - 1])
            {
                tau *= 0.5;
                goto tryAgain;
            }

            dN += (Nr[i] - 1.0) * V[i];

            // Detect sign change -> new partition boundary
            if (dV[i] * dN < 0.0)
            {
                i0[Ni0] = i;
                Ni0++;
            }
        }

        // Last tumor cell (with variable width hNN)
        dV[NN] = dN;

        dN += (Nr[NN] - 1.0) * (R_NN_h3 - R_NN3);

        if (dV[NN] * dN <= 0.0)
        {
            i0[Ni0] = NN;
            Ni0++;
        }

        hNNold = hNN;

        // Compute new radius of tumor domain from volume conservation
        double Rcur_old = R_NN + hNNold;     // OLD boundary before update
		double Rcur_old3 = Rcur_old * Rcur_old * Rcur_old;
		
		double arg = Rcur_old3 + dN;
		RNnew = (arg <= 0.0 ? 0.0 : fast_cbrt(arg));

        hNN = RNnew - R_NN;

		if (!RTflag && !NPflag && !ATflag) // without RT, NP and AT n=1 is kept within tumor, and all receptors are free
		{            
		    // If boundary exceeds one grid cell, add a new one
            while (hNN > dh)
            {
                NN  = NN + 1;
                n1[NN] = n1[NN - 1];  // copy boundary value
                ff1[NN] = ff1[NN - 1]; 
                fu1[NN] = fu1[NN - 1]; 
                hNN   = hNN - dh;
            }
            
			for (int i = 0; i <= NN; i++)
           		n1[i] = 1.;
		}
		else
		{
	        // Stability check on boundary movement
	        double dh_diff = hNN - hNNold;
	        if (dh_diff < 0.0) dh_diff = -dh_diff;
	        if (dh_diff > dh)
	        {
	            tau *= 0.5;
	            goto tryAgain;
	        }
	
	        // Determine direction of volume redistribution
	        // negative -> shrinkage starting from center
	        if (dV[1] < 0.0)
	            sign = -1;
	        else
	            sign = 1;
	            	
	        if (NPflag || ATflag)
				for (int i = 0; i <= NN; ++i)
			    {
			        double X = 0.0;
			        if (Nr[i] > 0.0)
			            X = nr[i] / Nr[i]; 
			
			        phifr[i] = ffr[i] * X;   // n * f_f
			        phiur[i] = fur[i] * X;   // n * f_u
			    }
			    
	        // --- Redistribution inside all subregions defined by i0[] ---
	        for (int ii = 1; ii < Ni0; ii++)
	        {
	            if (sign == -1)
	            {
	                // Block with mass moving outward (shrinkage near center)
	                for (int i = i0[ii - 1] + 1; i < i0[ii]; i++)
	                {
	                    n1[i] = ((nr[i] / Nr[i]) * (V[i] + dV[i + 1])
	                             - (nr[i + 1] / Nr[i + 1]) * dV[i + 1]) / V[i];
	                    if (NPflag || ATflag)
	                    {
		                    ff1[i] = ((phifr[i] * (V[i] + dV[i + 1]) - phifr[i+1] * dV[i+1]) / V[i]) / n1[i];
							fu1[i] = ((phiur[i] * (V[i] + dV[i + 1]) - phiur[i+1] * dV[i+1]) / V[i]) / n1[i];
	                	}
	                }
	
	                n1[i0[ii]] = nr[i0[ii]] / Nr[i0[ii]];
	                if (NPflag || ATflag)
	                {
		                ff1[i0[ii]] = phifr[i0[ii]]/n1[i0[ii]];
						fu1[i0[ii]] = phiur[i0[ii]]/n1[i0[ii]];
					}
	
	                sign = -sign;
	            }
	            else
	            {
	                // Block with inward redistribution (expansion)
	                if (ii == 1)
	                {
	                    n1[0] = nr[0] / Nr[0];
	                    if (NPflag || ATflag)
	              		{
	             	  		ff1[0] = phifr[0] / n1[0];
							fu1[0] = phiur[0] / n1[0];
						}
	
	                    for (int i = i0[ii - 1] + 2; i < i0[ii]; i++)
	                    {
	                        n1[i] = ((nr[i] / Nr[i]) * (V[i] - dV[i])
	                                 + (nr[i - 1] / Nr[i - 1]) * dV[i]) / V[i];
		                    if (NPflag || ATflag)
		              		{
	                        	ff1[i] = ((phifr[i] * (V[i] - dV[i]) + phifr[i-1] * dV[i]) / V[i])/n1[i];
								fu1[i] = ((phiur[i] * (V[i] - dV[i]) + phiur[i-1] * dV[i]) / V[i])/n1[i];
							}
	                    }
	                }
	                else
	                {
	                    for (int i = i0[ii - 1] + 1; i < i0[ii]; i++)
	                    {
	                        n1[i] = ((nr[i] / Nr[i]) * (V[i] - dV[i])
	                                 + (nr[i - 1] / Nr[i - 1]) * dV[i]) / V[i];
		                    if (NPflag || ATflag)
		              		{
	                        	ff1[i] = ((phifr[i] * (V[i] - dV[i]) + phifr[i-1] * dV[i]) / V[i])/n1[i];
								fu1[i] = ((phiur[i] * (V[i] - dV[i]) + phiur[i-1] * dV[i]) / V[i])/n1[i];
							}
	                    }
	                }
	
	                n1[i0[ii]] =
	                    ((nr[i0[ii]] / Nr[i0[ii]])
	                        * (V[i0[ii]] - dV[i0[ii]] + dV[i0[ii] + 1])
	                     + (nr[i0[ii] - 1] / Nr[i0[ii] - 1]) * dV[i0[ii]]
	                     - (nr[i0[ii] + 1] / Nr[i0[ii] + 1]) * dV[i0[ii] + 1])
	                    / V[i0[ii]];
	                
                    if (NPflag || ATflag)
              		{    
		                ff1[i0[ii]] = ((phifr[i0[ii]] * (V[i0[ii]] - dV[i0[ii]] + dV[i0[ii] + 1]) 
		                     + phifr[i0[ii] - 1] * dV[i0[ii]]
		                     - phifr[i0[ii] + 1] * dV[i0[ii] + 1])
		                    / V[i0[ii]])/n1[i0[ii]];
		                    
		                fu1[i0[ii]] = ((phiur[i0[ii]] * (V[i0[ii]] - dV[i0[ii]] + dV[i0[ii] + 1]) 
		                     + phiur[i0[ii] - 1] * dV[i0[ii]]
		                     - phiur[i0[ii] + 1] * dV[i0[ii] + 1])
		                    / V[i0[ii]])/n1[i0[ii]];	
		            }
	                    
	                sign = -sign;
	            }
	        }
	
	        // --- Final region near the boundary that contains hNN ---
	        if (hNN >= hNNold)   // tumor is expanding
	        {
	            if (Ni0 == 1)
	            {
	                n1[0] = nr[0] / Nr[0];
                    if (NPflag || ATflag)
              		{
                   		ff1[0] = phifr[0] / n1[0];
						fu1[0] = phiur[0] / n1[0];
					}
	
	                for (int i = i0[Ni0 - 1] + 2; i <= NN - 1; i++)
	                {
	                    // Standard outward redistribution
	                    n1[i] = ((nr[i] / Nr[i]) * (V[i] - dV[i])
	                             + (nr[i - 1] / Nr[i - 1]) * dV[i]) / V[i];	                             
	                    if (NPflag || ATflag)
	              		{
		                    ff1[i] = ((phifr[i] * (V[i] - dV[i]) + phifr[i-1] * dV[i]) / V[i]) / n1[i];
							fu1[i] = ((phiur[i] * (V[i] - dV[i]) + phiur[i-1] * dV[i]) / V[i]) / n1[i];
						}	
	                }
	            }
	            else
	            {
	                for (int i = i0[Ni0 - 1] + 1; i <= NN - 1; i++)
	                {
	                    n1[i] = ((nr[i] / Nr[i]) * (V[i] - dV[i])
	                             + (nr[i - 1] / Nr[i - 1]) * dV[i]) / V[i];
	                    if (NPflag || ATflag)
	              		{
		                    ff1[i] = ((phifr[i] * (V[i] - dV[i]) + phifr[i-1] * dV[i]) / V[i]) / n1[i];
							fu1[i] = ((phiur[i] * (V[i] - dV[i]) + phiur[i-1] * dV[i]) / V[i]) / n1[i];
						}
	                }
	            }
	
	            Vvar[NN] = R_NN_h3 - R_NN3;
	
	            if (Vvar[NN] == 0.0)
	            {
	                n1[NN] = nr[NN] / Nr[NN];
                    if (NPflag || ATflag)
              		{
	                    ff1[NN] = phifr[NN] / n1[NN];
						fu1[NN] = phiur[NN] / n1[NN];
					}
	            }
	            else
	            {
	                n1[NN] = ((nr[NN] / Nr[NN]) * (Vvar[NN] - dV[NN])
	                          + (nr[NN - 1] / Nr[NN - 1]) * dV[NN])
	                         / Vvar[NN];        
                    if (NPflag || ATflag)
              		{
		                ff1[NN] = ((phifr[NN] * (Vvar[NN] - dV[NN])
		                          + phifr[NN - 1] * dV[NN])
		                         / Vvar[NN]) / n1[NN];
		                fu1[NN] = ((phiur[NN] * (Vvar[NN] - dV[NN])
		                          + phiur[NN - 1] * dV[NN])
		                         / Vvar[NN]) / n1[NN];
		           	}
	            }
	
	            // If boundary exceeds one grid cell, add a new one
	            if (hNN > dh)
	            {
	                NN  = NN + 1;
	                n1[NN] = n1[NN - 1];  // copy boundary value
	                ff1[NN] = ff1[NN - 1]; 
                	fu1[NN] = fu1[NN - 1]; 
	                hNN   = hNN - dh;
	            }
	        }
	        else   // hNN < hNNold: tumor is shrinking
	        {
	            if (hNN >= 0.0)
	            {
	                // Shrinkage without removing a whole grid cell
	                for (int i = i0[Ni0 - 1] + 1; i < NN; i++)
	                {
	                    n1[i] = ((nr[i] / Nr[i]) * (V[i] + dV[i + 1])
	                             - (nr[i + 1] / Nr[i + 1]) * dV[i + 1]) / V[i];
	                    if (NPflag || ATflag)
	              		{
		                    ff1[i] = ((phifr[i] * (V[i] + dV[i + 1])
		                             - (phifr[i + 1] * dV[i + 1])) / V[i]) / n1[i];
		                    fu1[i] = ((phiur[i] * (V[i] + dV[i + 1])
		                             - (phiur[i + 1] * dV[i + 1])) / V[i]) / n1[i];
	                   }
	                }
	
	                n1[NN] = nr[NN] / Nr[NN];
                    if (NPflag || ATflag)
              		{
		                ff1[NN] = phifr[NN] / n1[NN];
		                fu1[NN] = phiur[NN] / n1[NN];
	            	}
	            }
	            else
	            {
	                // Full removal of the last grid cell
	                for (int i = i0[Ni0 - 1] + 1; i < NN - 1; i++)
	                {
	                    n1[i] = ((nr[i] / Nr[i]) * (V[i] + dV[i + 1])
	                             - (nr[i + 1] / Nr[i + 1]) * dV[i + 1]) / V[i];	                    
	                    if (NPflag || ATflag)
	              		{
		                    ff1[i] = ((phifr[i] * (V[i] + dV[i + 1])
		                             - (phifr[i + 1] * dV[i + 1])) / V[i]) / n1[i];
		                    fu1[i] = ((phiur[i] * (V[i] + dV[i + 1])
		                             - (phiur[i + 1] * dV[i + 1])) / V[i]) / n1[i];
	                	}
	                }
	
	                // Volume of new last shell [NN-1, NN-1+hNN]
	                double R_NNm1    = (NN - 1) * dh;
	                double R_NNm13   = R_NNm1 * R_NNm1 * R_NNm1;
	                double R_last_h3 = R_NN_h3;  // same as (NN*dh + hNN)^3
	
	                Vvar[NN - 1] = R_last_h3 - R_NNm13;
	
	                n1[NN - 1] =
	                    ((nr[NN - 1] / Nr[NN - 1]) * (Vvar[NN - 1] + dV[NN])
	                     - (nr[NN] / Nr[NN]) * dV[NN])
	                    / Vvar[NN - 1];	     
                    if (NPflag || ATflag)
              		{           
		                ff1[NN - 1] =
		                    ((phifr[NN - 1] * (Vvar[NN - 1] + dV[NN])
		                     - phifr[NN] * dV[NN])
		                    / Vvar[NN - 1]) / n1[NN - 1];              
		                fu1[NN - 1] =
		                    ((phiur[NN - 1] * (Vvar[NN - 1] + dV[NN])
		                     - phiur[NN] * dV[NN])
		                    / Vvar[NN - 1]) / n1[NN - 1];
	                }
	
	                n1[NN] = 0.0;
	                ff1[NN] = 0.0;
                	fu1[NN] = 0.0;
	
	                NN  = NN - 1;
	                hNN = dh + hNN;
	            }
	        }   
	        
			for (int i = 0; i <= NN; ++i)
			{
			    bool bad = (n1[i] < 1e-12) ||
			               (!std::isfinite(n1[i])) ||
			               (!std::isfinite(ff1[i])) ||
			               (!std::isfinite(fu1[i]));
			
			    if (bad)
			    {
			        n1[i] = CLAMP01(n1[i]);
			        ff1[i] = ff0[i];
			        fu1[i] = fu0[i];
			    }
			    else
			    {
			        n1[i] = CLAMP01(n1[i]);
			        ff1[i] = CLAMP01(ff1[i]);
			        fu1[i] = CLAMP01(fu1[i]);
			    }
			}
			
			// Reset receptors in normal tissue
			for (int i = NN+1; i <= NH; i++) 
			{
			    ff1[i] = 0.0;
			    fu1[i] = 0.0;
			}
		}
    }

    // --- 3) Update total tissue radius (healthy tissue relocation) ---

    double Rtum = NN * dh + hNN;
    if (Rtum < 0.0) Rtum = 0.0;

    // Volume conservation: R_tissue^3 = R0^3 + R_tumor^3
    double R0T3  = R0T * R0T * R0T;
    double Rtum3 = Rtum * Rtum * Rtum;
    double arg = R0T3 + Rtum3;
	RHnew = (arg <= 0.0) ? 0.0 : fast_cbrt(arg);

    NH  = (int)(RHnew / dh);
    hNH = RHnew - 1.0 * NH * dh;
    
    
}

//********* QUASI-STATIONARY APPROXIMATIONS FOR GLUCOSE

void Glucose_qs()
{
    // Case NN == 0: tumor is extremely small and does not affect glucose distribution
    if (NN == 0)
    {
        for (int i = 0; i <= NH; i++)
            g1[i] = gstat;
    }
    else
    {
        // --- 1) Initial preparation ---
        // Convert g0 to G0 = g0 * r; radii (r) are precomputed in Rmid[]
        for (int i = 0; i <= NH; i++)
            G0[i] = g0[i] * Rmid[i];

        norm_g = 1.0;

        // Precompute inverses for faster arithmetic
        const double inv_kg     = 1.0 / kg;
        const double Pg_over_kg = Pg * inv_kg;
        const double Q_over_kg  = Q * inv_kg;

        // --- 2) Build Fg[i] and alpha_g[i] outside the iteration loop ---

        // Inside the tumor (except last tumor cell NN)
        for (int i = 1; i <= NN - 1; i++)
        {
            Fg[i]      = 0.0;
            alpha_g[i] = Q_over_kg * n0[i];
        }

        // Geometry at the tumor boundary (cell NN and NN+1)
        const double R_NN    = NN * dh;
        const double R_NNp1  = (NN + 1) * dh;
        const double R_NNh   = R_NN + hNN;

        const double R_NN3   = R_NN   * R_NN   * R_NN;
        const double R_NNp13 = R_NNp1 * R_NNp1 * R_NNp1;
        const double R_NNh3  = R_NNh  * R_NNh  * R_NNh;

        const double full_shell = R_NNp13 - R_NN3;   // Full volume of shell [NN, NN+1]
        const double tum_shell  = R_NNh3  - R_NN3;   // Portion filled by tumor
        const double out_shell  = R_NNp13 - R_NNh3;  // Outer portion

        const double ratio_tum = tum_shell / full_shell;
        const double ratio_out = out_shell / full_shell;

        // Boundary flux at NN
        Fg[NN]      = Pg_over_kg * ratio_out;
        alpha_g[NN] = Q_over_kg  * n0[NN] * ratio_tum;

        // Outside the tumor (NN+1..NH-1)
        for (int i = NN + 1; i <= NH - 1; i++)
        {
            Fg[i]      = Pg_over_kg * Rmid[i];
            alpha_g[i] = 0.0;
        }

        // --- 3) Iterative tridiagonal solver ---

        kqs = 1;

        // Right boundary factor does not depend on iteration
        hirightg = ((double)NH + 0.5) / ((double)NH - 0.5);

        while ((norm_g > norm_min) && (kqs < k_crit))
        {
            // 3.1. Assemble the main diagonal bprg[i]

            // Inside the tumor
            for (int i = 1; i <= NN - 1; i++)
            {
                double g_loc = G0[i] * Rmid_inv[i];   // G0[i] / r_i
                bprg[i] = 2.0 + alpha_g[i] / (g_loc + gst);
            }

            // Tumor boundary (i = NN)
            {
                double g_loc_NN = G0[NN] * Rmid_inv[NN];
                bprg[NN] = 2.0
                         + Pg_over_kg * ratio_out
                         + alpha_g[NN] / (g_loc_NN + gst);
            }

            // Outside tumor
            for (int i = NN + 1; i <= NH - 1; i++)
            {
                double g_loc = G0[i] * Rmid_inv[i];
                bprg[i] = 2.0 + Pg_over_kg + alpha_g[i] / (g_loc + gst);
            }

            // 3.2. Solve the tridiagonal system
            TMA(aprg, bprg, cprg, Fg, progg,
                muleftg, hileftg, murightg, hirightg);

            // 3.3. Compute L norm and update G0
            norm_g = 0.0;

            for (int i = 0; i <= NH; i++)
            {
                G1[i] = progg[i];

                double diff = G0[i] - G1[i];
                if (diff < 0.0) diff = -diff;
                if (diff > norm_g)
                    norm_g = diff;

                G0[i] = G1[i];
            }

            kqs++;
        }

        // --- 4) Convert back to g0, g1 = G / r ---
        for (int i = 0; i <= NH; i++)
        {
            g0[i] = G0[i] * Rmid_inv[i];
            g1[i] = G1[i] * Rmid_inv[i];
        }
    }
}

//***********************************************************************************************************************************************************
//********************************* DIFFUSION ***************************************************************************************************************
//***********************************************************************************************************************************************************

void Diffusion_NP()
{
	for (int i = 0; i <= NH; i++)
	{
		U0[i] = u0[i] * (1. * i + 0.5) * dh; // for easy solution in the spherical symmetrical case
		UR[i] = ur[i] * (1. * i + 0.5) * dh;
	}

	for (int i = 1; i <= NH - 1; i++)
		Fu[i] = (U0[i + 1] - 2. * U0[i] + U0[i - 1]) + ku * UR[i];

	hirightu = (1. * NH + 0.5) / (1. * NH - 0.5);

	TMAc(apru, bpru, cpru, Fu, progu, muleftu, hileftu, murightu, hirightu);

	for (int i = 0; i <= NH; i++)
		U1[i] = progu[i];

	for (int i = 0; i <= NH; i++)
		u1[i] = U1[i] / ((1. * i + 0.5) * dh);
}

void Diffusion_AT()
{
	for (int i = 0; i <= NH; i++)
	{
		A0[i] = a0[i] * (1. * i + 0.5) * dh; // for easy solution in spherical symmetrical case
		AR[i] = ar[i] * (1. * i + 0.5) * dh;
	}

	for (int i = 1; i <= NH - 1; i++)
		Fa[i] = (A0[i + 1] - 2. * A0[i] + A0[i - 1]) + ka * AR[i];

	hirighta = (1. * NH + 0.5) / (1. * NH - 0.5);

	TMAc(apra, bpra, cpra, Fa, proga, mulefta, hilefta, murighta, hirighta);

	for (int i = 0; i <= NH; i++)
		A1[i] = proga[i];

	for (int i = 0; i <= NH; i++)
		a1[i] = A1[i] / ((1. * i + 0.5) * dh);
}

//********* TRIDIAGONAL MATRIX ALGORITHM

void TMA(double apr, double* bpr, double cpr, double* F, double* prog,
	double muleft, double hileft, double muright, double hiright)
{
	pp[1] = hileft;
	qq[1] = muleft;

	for (int i = 1; i <= NH - 1; i++)
	{
		pp[i + 1] = cpr / (bpr[i] - pp[i] * apr);
		qq[i + 1] = (apr * qq[i] + F[i]) / (bpr[i] - pp[i] * apr);
	}

	prog[NH] = (muright + hiright * qq[NH]) / (1. - pp[NH] * hiright);

	for (int i = NH - 1; i >= 0; i--)
		prog[i] = pp[i + 1] * prog[i + 1] + qq[i + 1];
}

void TMAc(double apr, double bpr, double cpr, double* F, double* prog,
	double muleft, double hileft, double muright, double hiright)
{
	pp[1] = hileft;
	qq[1] = muleft;

	for (int i = 1; i <= NH - 1; i++)
	{
		pp[i + 1] = cpr / (bpr - pp[i] * apr);
		qq[i + 1] = (apr * qq[i] + F[i]) / (bpr - pp[i] * apr);
	}

	prog[NH] = (muright + hiright * qq[NH]) / (1. - pp[NH] * hiright);

	for (int i = NH - 1; i >= 0; i--)
		prog[i] = pp[i + 1] * prog[i + 1] + qq[i + 1];
}

//***********************************************************************************************************************************************************
//********************************* RADIOTHERAPY *******************************************************************************************************
//***********************************************************************************************************************************************************

void PrepareRT()
{
    for (int i = 0; i <= NN; i++)
    {
        // ---- Effective dose ----
        if (!NPlimmode)
        {
            Deff[i] = D * (1.0 + Ku * (rho * fu0[i] * n0[i] + u0[i]));
        }
        else // limiting NP mode: fu = 1, no u
        {
            Deff[i] = D * (1.0 + Ku * (rho * n0[i]));
        }

        // ---- Proliferation-dependent alpha ----
        alpha_eff[i] = alpha0 + (alphap - alpha0) * g0[i] / (g0[i] + gst);

        // ---- LQ exponent ----
        RTcoef[i] = Deff[i] * alpha_eff[i] * (1.0 + betaalpha * Deff[i]);

        // ---- Survival ----
        nRTsurv[i] = exp(- RTcoef[i]);
    }
}

void AssessRT()
{
	SURV = 0.;

	for (int i = 0; i <= NN - 1; i++)
		SURV = SURV + V[i] * n0[i] * nRTsurv[i];

	SURV = SURV + ((NN*dh + hNN)*(NN*dh + hNN)*(NN*dh + hNN) - NN*dh*NN*dh*NN*dh) * n0[NN] * nRTsurv[NN];

	SURV = 4. * PI * 300. * SURV / 3.;
	
	if (SURV < SURVmin)
	{
		SURVmin = SURV;
		tSURVmin = t - tu;
	}
}

void PerformRT()
{	
    // ---- Apply radiation damage ----
    for (int i = 0; i <= NN; i++)
    {
        n0[i] *= nRTsurv[i];
        n1[i]  = n0[i];
    }
}

//***********************************************************************************************************************************************************
//********************************* MAIN CYCLE ENDING *******************************************************************************************************
//***********************************************************************************************************************************************************

void Cycle_Ending()
{
	for (int i = 0; i <= NH; i++)
	{
		n0[i] = n1[i];
		g0[i] = g1[i];
		u0[i] = u1[i];
		ff0[i] = ff1[i];
		fu0[i] = fu1[i];
		a0[i] = a1[i];
	}
}

//***********************************************************************************************************************************************************
//********************************* PROGRAM ENDING **********************************************************************************************************
//***********************************************************************************************************************************************************

void End()
{
	if (NTC<0.01)
		survival = Tsurv/24.;
	else
		survival = (t - survival)/24.;
	
    char survname[256];
    sprintf(survname, "Survival.txt");

    FILE* out = fopen(survname, "wt");
    if (!out)
    {
        fprintf(stderr, "Cannot open survival file %s\n", survname);
        return;
    }

    fprintf(out, "%.3f", survival);

    fclose(out);
    
    if (RTmode && tNPRT == 999999999.0)
    {
		char survminname[256];
	    sprintf(survminname, "MinSurvCells.txt");
	
	    FILE* outtwo = fopen(survminname, "wt");
	    if (!outtwo)
	    {
	        fprintf(stderr, "Cannot open file for minimal number of cells that would survive %s\n", survminname);
	        return;
	    }
	
	    fprintf(outtwo, "%.3f\t%.3e", tSURVmin/24., SURVmin);
	
	    fclose(outtwo);
	}
	
	if (clock() / CLOCKS_PER_SEC >= 3600)
		printf("\n%s%ld%s%ld%s%ld%s", "The end. Total time: ", (clock() / CLOCKS_PER_SEC) / 3600, " h ", (clock() / CLOCKS_PER_SEC) / 60 - 60 * ((clock() / CLOCKS_PER_SEC) / 3600), " min ", (clock() / CLOCKS_PER_SEC) - 3600 * ((clock() / CLOCKS_PER_SEC) / 3600) - 60 * ((clock() / CLOCKS_PER_SEC) / 60 - 60 * ((clock() / CLOCKS_PER_SEC) / 3600)), " s.");
	else if (clock() / CLOCKS_PER_SEC >= 60)
		printf("\n%s%ld%s%ld%s", "The end. Total time: ", (clock() / CLOCKS_PER_SEC) / 60 - 60 * ((clock() / CLOCKS_PER_SEC) / 3600), " min ", (clock() / CLOCKS_PER_SEC) - 3600 * ((clock() / CLOCKS_PER_SEC) / 3600) - 60 * ((clock() / CLOCKS_PER_SEC) / 60 - 60 * ((clock() / CLOCKS_PER_SEC) / 3600)), " s.");
	else
		printf("\n%s%ld%s", "The end. Total time: ", (clock() / CLOCKS_PER_SEC) - 3600 * ((clock() / CLOCKS_PER_SEC) / 3600) - 60 * ((clock() / CLOCKS_PER_SEC) / 60 - 60 * ((clock() / CLOCKS_PER_SEC) / 3600)), " s.");
	//_getch();
	exit(0);
}

//********************************************************
//********* THE END
//********************************************************
```
